# Open chromatin analysis in *Trypanosoma cruzi* life forms highlights critical differences in genomic compartments and developmental regulation at tDNA loci

**DOI:** 10.1101/2021.10.26.465934

**Authors:** Alex Ranieri Jerônimo Lima, Saloe Bispo Poubel, Juliana Nunes Rosón, Loyze Paola Oliveira de Lima, Héllida Marina Costa-Silva, Herbert Guimarães de Sousa Silva, Camila Silva Gonçalves, Pedro A. F. Galante, Fabiola Holetz, Maria Cristina Machado M. Motta, Ariel M. Silber, M. Carolina Elias, Julia Pinheiro Chagas da Cunha

## Abstract

**Background:** Genomic organization and gene expression regulation in trypanosomes are remarkable because protein-coding genes are organized into codirectional gene clusters with unrelated functions. Moreover, there is no dedicated promoter for each gene, resulting in polycistronic gene transcription, with posttranscriptional control playing a major role. Nonetheless, these parasites harbor epigenetic modifications at critical regulatory genome features that dynamically change among parasite stages, which are not fully understood.

**Results:** Here, we investigated the impact of chromatin changes in a scenario commanded by posttranscriptional control exploring the parasite *Trypanosoma cruzi* and its differentiation program using genome-wide approaches supported by transmission electron microscopy. The integration of FAIRE and MNase-seq data, two complementary epigenomic approaches, enabled us to identify differences in *T. cruzi* genome compartments, putative transcriptional start regions and virulence factors. In addition, we also detected developmental chromatin regulation at tRNA loci (tDNA), which seems to be linked to the translation regulatory mechanism required for parasite differentiation. Strikingly, a positive correlation was observed between active chromatin and steady-state transcription levels.

**Conclusion:** Taken together, our results indicate that chromatin changes reflect the unusual gene expression regulation of trypanosomes and the differences among parasite developmental stages, even in the context of a lack of canonical transcriptional control of protein-coding genes.

## Background

Chromatin is the stage where gene expression and epigenetic regulation occur. The epigenome of a cell consists of a series of heritable chromatin modifications that orchestrate and mirror essential aspects of gene regulation. For example, in general, the promoter regions of highly expressed genes have an open chromatin state and are enriched in active histone marks such as H3K4me. In contrast, genomic regions with lower transcription rates are present in a more compact chromatin state and enriched in histone marks such H3K9me3 and H3K27me3 (1, 2). Epigenomic changes can be explored using next-generation sequencing coupled with traditional approaches to detect DNA regulatory sequences. The FAIRE (formaldehyde-assisted isolation of regulatory elements)-seq technique is one of these approaches that uses formaldehyde to cross-link proteins, mainly histones, to DNA (3). This results in an enrichment of genomic regions associated with regulatory activity, previously detected by DNAse I and active chromatin marks, such as H3K4me and RNA Pol II (4). Promoter regions of highly expressed genes are more enriched in FAIRE than poorly transcribed regions, indicating that FAIRE-enriched sequences are associated with active genomic regions (5).

Trypanosomatids are early-divergent eukaryotes that present some unique molecular characteristics, such as a hypermodified thymine base, known as base J, intronless genes, polycistronic transcription of functionally unrelated genes, mRNA maturation by trans-splicing, and posttranscriptional regulation of gene expression (6). RNA Pol II transcription starts mainly at the intergenic regions between two polycistronic transcription units (PTUs) of divergent transcription (dSSR – divergent strand switch region) and ends between two convergent PTUs (cSSR – convergent strand switch region) or at regions transcribed by other polymerases (Clayton 2019). Although no gene-specific promoters have been described, transcription starts at dispersed promoter regions rich in GT (7) and decorated by epigenetic marks and histone variants (H2A. Z and H2B. V) (Respuela, Ferella, and Rada-Iglesias 2008; Siegel et al. 2009; Wright, Siegel, and Cross 2010). Transcription termination regions (TTRs) also contain epigenetic marks, such as histone variant H3. V and H4. V, which favor ending RNA Pol II transcription (10, 11). In addition, dSSRs and cSSRs were shown to be enriched in base J (12).

*Trypanosoma cruzi,* the etiologic agent of Chagas disease, alternates between a vertebrate host and a triatomine vector (13). In addition to the differences in cell shape, organization, and metabolism, *T. cruzi* life forms are also distinguished by the capacity of cell replication (epimastigotes and amastigotes) and cell infection (tissue culture trypomastigotes – TCT; and metacyclic trypomastigotes - MTs) (14–16). Profound changes in global transcription are found during the differentiation of epimastigotes into MTs and between epimastigotes and TCTs (17). Evaluation of transcription rate dynamics during metacyclogenesis indicates that the decrease in transcription rates only occurs in fully differentiated MTs (18). Differentiation to MTs is followed by a global downregulation of the translatome except by a translational upregulation of virulence factors, such as TS and GP63, in MTs (19). The phosphorylation status of the translation factors eIF2α and eIF5a seems to be a key mechanism controlling translation and, therefore, *T. cruzi* metacyclogenesis (20, 21).

Nucleus and chromatin also change throughout the parasite life cycle. In epimastigotes and amastigotes, the nucleus is spherical, containing a large nucleolus and high amounts of euchromatin, which is considered transcriptionally active. In contrast, the trypomastigote forms (TCTs and MTs) present an elongated nucleus, absence of nucleolus, and abundant heterochromatin, which is dispersed in the nucleoplasm (15,17,18). Changes in nuclear morphology start at the beginning of metacyclogenesis, while heterochromatin spreads progressively along with the nucleus in intermediate forms (15). The mechanisms that drive and control these ultrastructural alterations during metacyclogenesis are not fully understood.

The genome of this parasite was proposed to be compartmentalized in two regions that differ in gene and GC content: the core compartment (composed of conserved and hypothetical conserved genes) and the disruptive compartment (containing the multigenic families trans-sialidase – TS, mucin-associated surface proteins – MASP, and mucins). GP63, dispersed gene family 1 (DGF-1), and retrotransposon hot spot (RHS) multigene families can occur in both compartments (22). Multigene family members such as TS, mucins, MASP, and GP63 code for virulence factors present on the parasite surface and are upregulated in infective forms (23).

In recent years, our group has demonstrated that changes in epigenetic marks, such as histone posttranslational modifications (PTMs), histone variant deposition, and nucleosome occupancy, change among *T. cruzi* life forms and follow parasite differentiation (24–26). As *T. cruzi* differentiation is followed by a global decrease in transcription, translation, and alterations in nuclear morphology in a background of transcriptional regulation absence of protein-coding genes, we comprehensively investigated the impact of chromatin changes in a scenario commanded by posttranscriptional control. By integrating the open chromatin mapping obtained by FAIRE-seq with public transcriptomic and nucleosome mapping datasets, we identified important differences in *T. cruzi* genome compartments and a developmental change at tRNA loci (tDNA). Of note, the latter may be associated with the regulatory mechanism associated with translation control required for parasite differentiation. In addition, to shed light on the mechanisms related to the differentiation of infective forms, our data can disentangle the impact of chromatin remodeling in a scenario commanded mainly by posttranscriptional control.

## Results

### Core and disruptive genomic compartments of *T. cruzi* have a different open chromatin profile

To investigate whether the lack of transcriptional regulation of protein-coding genes would be reflected in the open chromatin profile of *T. cruzi*, we performed a FAIRE-seq genome-wide approach. We rescued these regions from the genome of epimastigotes under exponential proliferation. Biological duplicates were processed together with input control samples (reversed cross-link before DNA extraction) to account for any bias in genome sequencing and assembly. FAIRE-seq reads presented high duplication levels (66.2-72.1% in epimastigotes – **Figure S1 A**) and high Phred score (quality) values (>30 – **Figure S1 B**). After the mapping process (Figure S1 C), only mapped reads with mapping quality scores (MAPQs) above 10 were retained, resulting in approximately 116 million reads mapped (**Figure S1 D)**. Spearman correlation analysis (**Figure S2 A**) and PCA (**Figure S2 B**) using genome-wide read coverage of 50 bp windows for each replicate indicated a high correlation among them, showing good reproducibility and agreement among biological replicates. To avoid bias at repetitive regions, we removed multimapped reads, resulting in reduced (or zero) read coverage at repetitive regions of the genome (**Figure S3**). Most of the repetitive regions occurred in intergenic sections (92.75%) of the genome (**Supplementary Table S1**). Of note, strand switch regions (SSRs) were not considered intergenic regions; nevertheless, no repeat annotations were found there. Only 309 CDSs (5.65% of total repetitive regions, in bp) were located in annotated repetitive regions.

To obtain an overview of the open chromatin distribution in large genomic regions, we compared the data of epimastigote FAIRE-seq (E) and their corresponding control samples (C). Figure 1 shows the distribution of reads obtained from both samples on two contigs from Dm28c (PRFA1000011 – 594 kb and PRFA1000027 – 454 kb). Datasets were normalized to reads per genomic content (RPGC) to account for different sequencing depths. Visual inspection of RPGC levels indicates that the FAIRE sample has higher overall levels along the contigs with some clear enrichments when compared to the control sample. Ratio levels in log2 (E/C) indicated that most regions were enriched in FAIRE samples (**Figure 1A**). Repetitive regions are depleted in both FAIRE and control samples, likely due to the applied mapping and filtering steps (**Figure S3-4**).

**Figure 1.**
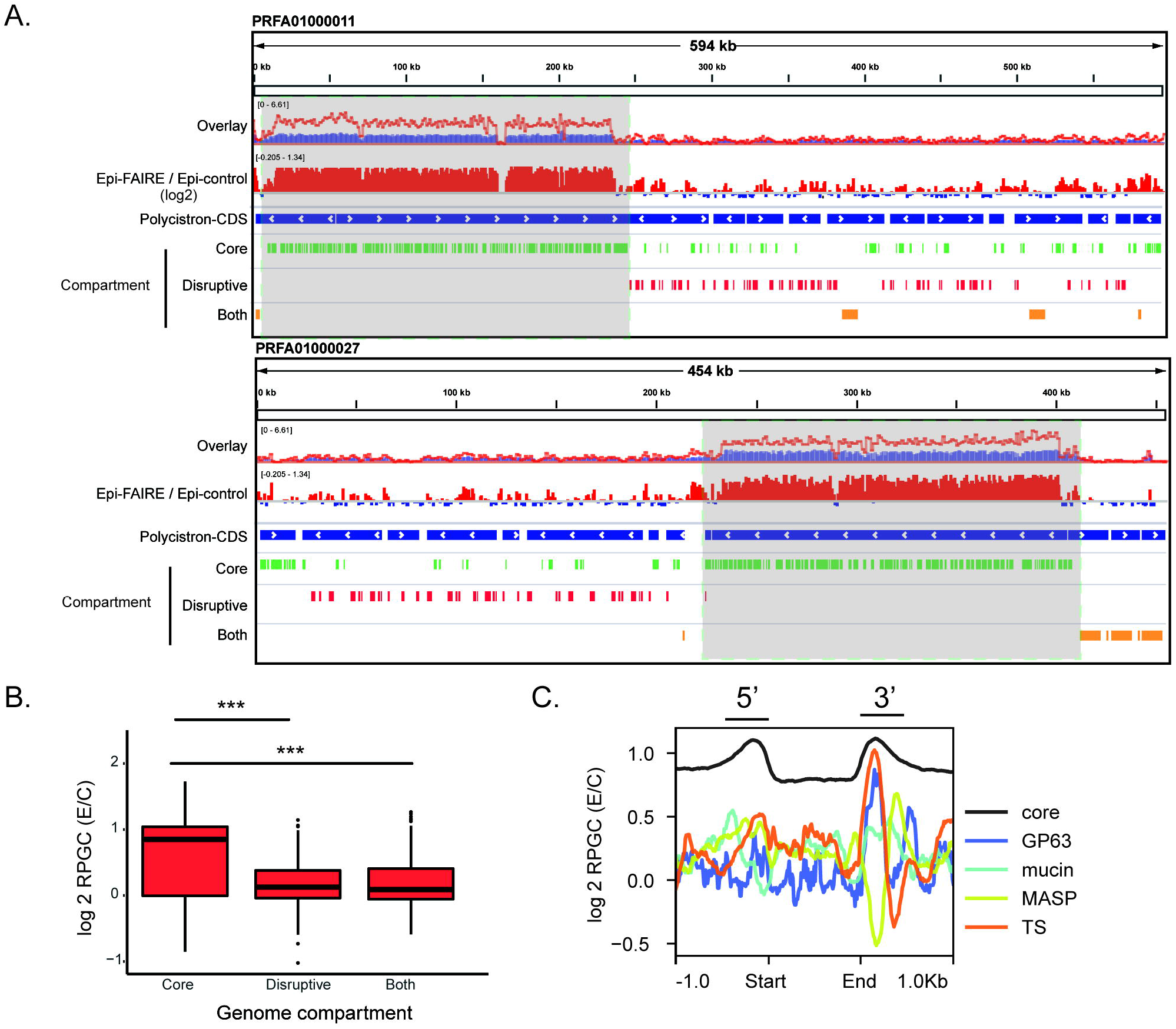
Genome-wide analysis of active chromatin in *T. cruzi*. (**A**) Overlay of the number of reads per genomic content (RPGC) in FAIRE-seq data for epimastigotes (red line) and its control (blue) on contigs PRFA01000011 and PRFA01000027. Track with the ratio FAIRE/control is depicted in the log2 ratio. Polycistrons are shown in blue tracks; arrows indicate the transcription direction; genes from the conserved compartment, disruptive compartment and both genome compartments are shown in green, red, and orange tracks, respectively. **B.** RPGC log2 ratio of normalized epimastigote FAIRE reads in different *T. cruzi* compartments. **C.** Active chromatin landscape in core genes (nonmultigenic family) and virulence factors 1 kb upstream and downstream of their respective start (ATG) and end (STOP codon) coding regions. (*** Wilcoxon-Mann–Whitney test with p-value = 0.001)

Regions covered by the core compartment (composed of conserved and hypothetical conserved genes, green tracks) had higher RPGC normalized counts than the disruptive compartment regions (red tracks) (**Figure 1A and B, Table S2-3**). Differences in RPGC counts between these two compartments were also observed in control samples; however, this difference was even more evident in FAIRE samples, which was reflected by higher levels of log2 ratio (E/C) in the core than in the disruptive compartment (**Figure 1 B**). Remarkably, removing multimapped reads (see methods - **Figure S4**) slightly affected this difference since the same criteria were used for experimental and control samples. Instead, this can be explained by the less open chromatin at the multigenic family in contrast to a more open chromatin profile in the nonmultifamily CDS.

Because the distribution of open chromatin was revealed to be distinct between genome compartments, we asked whether their landscapes, at a gene level, were also different. Strikingly, the landscape of open chromatin greatly differs among virulence factors (present in the disruptive compartment), among them, and among the remaining protein-coding genes. **Figure 1C** shows that at most core-gene members, the regions coding for their 5’ and 3’ UTRs are enriched in open chromatin, corroborating the impoverishment of nucleosomes seen previously by MNASE-seq (**Figure S5**). In contrast, no clear enrichment of open chromatin was found around the genomic regions coding for the 5’ UTR of virulence factors, while the regions coding for the 3’ UTR greatly differed among them: TS and GP63 exhibited an enrichment similar to other CDSs. At the same time, MASPs had a clear depletion followed by an enrichment a few base pairs downstream (**Figure 1C**). Nucleosome occupancy partially explains the active chromatin distribution around the virulence factor genes, as some regions (such as the upstream regions of mucins, GP63, DGF-1 and RHS) exhibit enrichment or decrease both in FAIRE and MNAse data (**Figure S5**).

### Open chromatin is enriched at divergent SSRs (dSSRs) and uniformly distributed along PTUs

In trypanosomatids, transcription of protein-coding genes initiates mainly at dSSRs and terminates at cSSRs; however, transcription might also start at some non- SSRs, mainly close to tRNA genes (27)(6, 28). This latter has not yet been described in *T. cruzi,* we inspected dSSRs as a proxy of transcription start sites, which resulted in enrichment of open chromatin compared to PTUs, as evidenced in **Figure 2A and 2B**. Combining a previous nucleosome mapping and occupancy profile (25) with the FAIRE-seq data along PTUs revealed complementary opposite landscape profiles: open chromatin regions mainly reflect nucleosome-depleted regions (**Figure 2A**).

**Figure 2.**
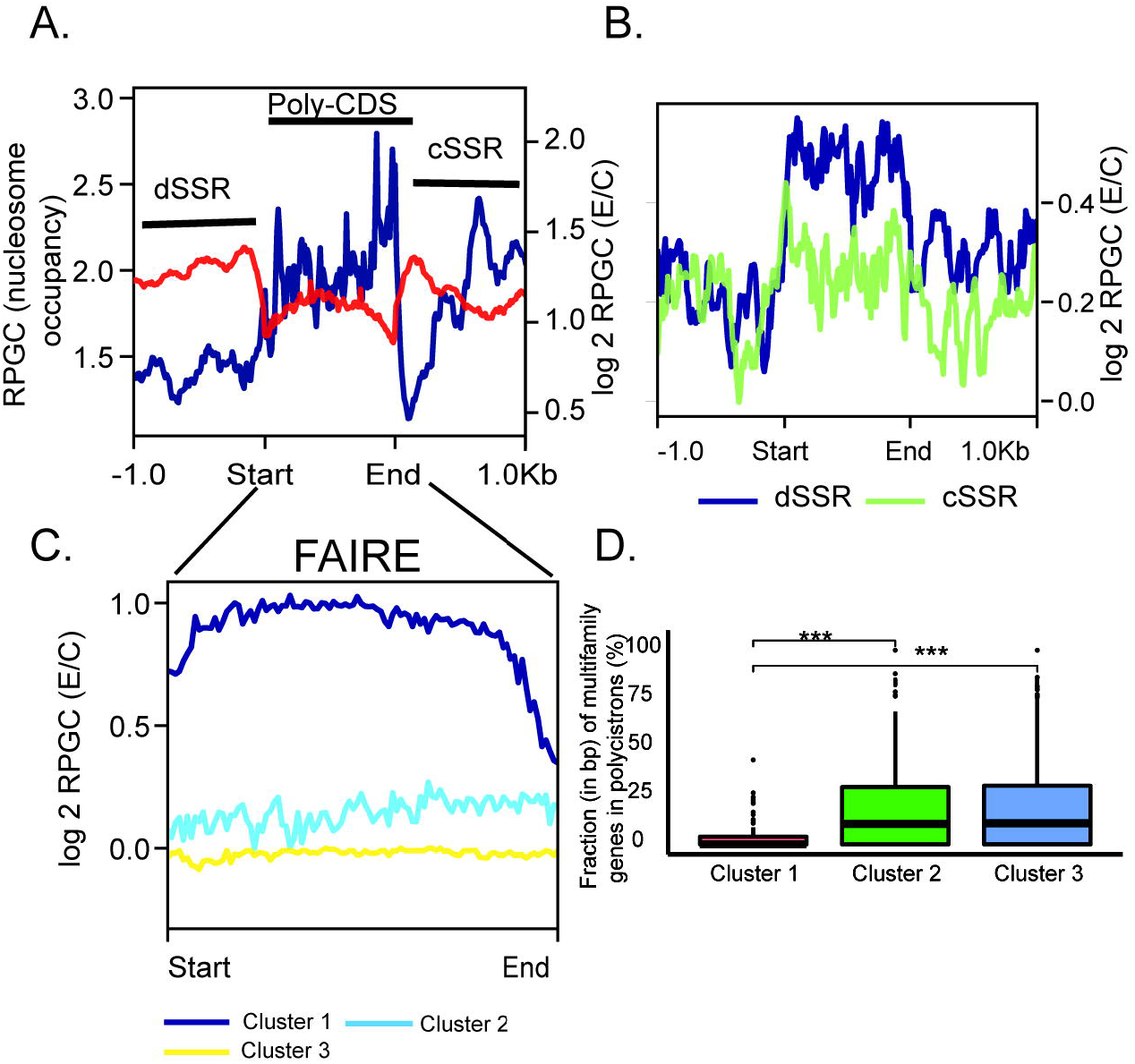
Open chromatin at a PTU context in epimastigotes. **A.** Superposition of FAIRE (red – right values on the y-axis) and MNase-seq (blue – left values on the y-axis) datasets. **B.** Log2 ratio of normalized epimastigote FAIRE reads in dSSR (blue) and cSSR (green). **C.** Hierarchical cluster analysis of FAIRE data depicted in **A** (Cluster 1: n = 227; Cluster 2: n = 412; Cluster 3: n = 258). In **A**, **B** and **C**, the first base of the feature is Start, while the last base is End. **D.** Percentage of bases from multifamily genes for each PTU according to hierarchical cluster analysis shown in **C**. (*** Wilcoxon-Mann-Whitney test with p-value = 0.001). E/C, ratio for epimastigotes and their respective controls.

PTU regions were hierarchically clustered into three groups based on their log2 (E/C) RPGC level (**Figure 2 C**). Clusters 2 and 3 are very similar, exhibiting a near flat pattern with low overall RPGC levels. In contrast, Cluster 1 showed higher overall levels of open chromatin, with a decrease at the edges, mainly at the PTU ending region. Clusters 2 and 3 contained significantly more genes (per bp) from the multigenic family than Cluster 1, which was enriched mainly from genes of the core compartment (**Figure 2D**).

### Levels and relative nuclear position of eu- and heterochromatin changes during metacyclogenesis

Many morphological changes are observed during the differentiation of epimastigotes to MTs, including nuclear elongation and kinetoplast repositioning to the parasite posterior end. Previously, it has been reported that the heterochromatin near the nuclear envelope in the epimastigote forms spreads progressively along with the nucleus in the intermediate forms, reaching a higher level of compaction in metacyclics in MTs (15). However, only in this work a systematic evaluation and 3D reconstruction of the two traditional chromatin classes, eu- and heterochromatin, was performed to check chromatin remodeling during parasite differentiation. The obtained results indicate that in *T. cruzi* epimastigotes, euchromatin resides in the central area, whereas heterochromatin is mainly distributed close to the nuclear envelope and surrounding the nucleolus (**Figure 3A** **– epimastigote**). The euchromatin volume is higher in epimastigote and intermediate I forms and decreases during differentiation to MT. Heterochromatin, in the epimastigote form, spreads throughout the nucleus, and as metacyclogenesis advances, its percentage increases and its location becomes increasingly peripheral. These results indicate that progressive chromatin remodeling occurs during parasite differentiation. It is worth observing that during metacyclogenesis, the nucleolus reduces its size, as it undergoes disassembly and dispersion throughout the nuclear matrix, which may be related to the decrease in ribosomal biogenesis (**Figure 3, Supplemental videos S1-4**).

**Figure 3.**
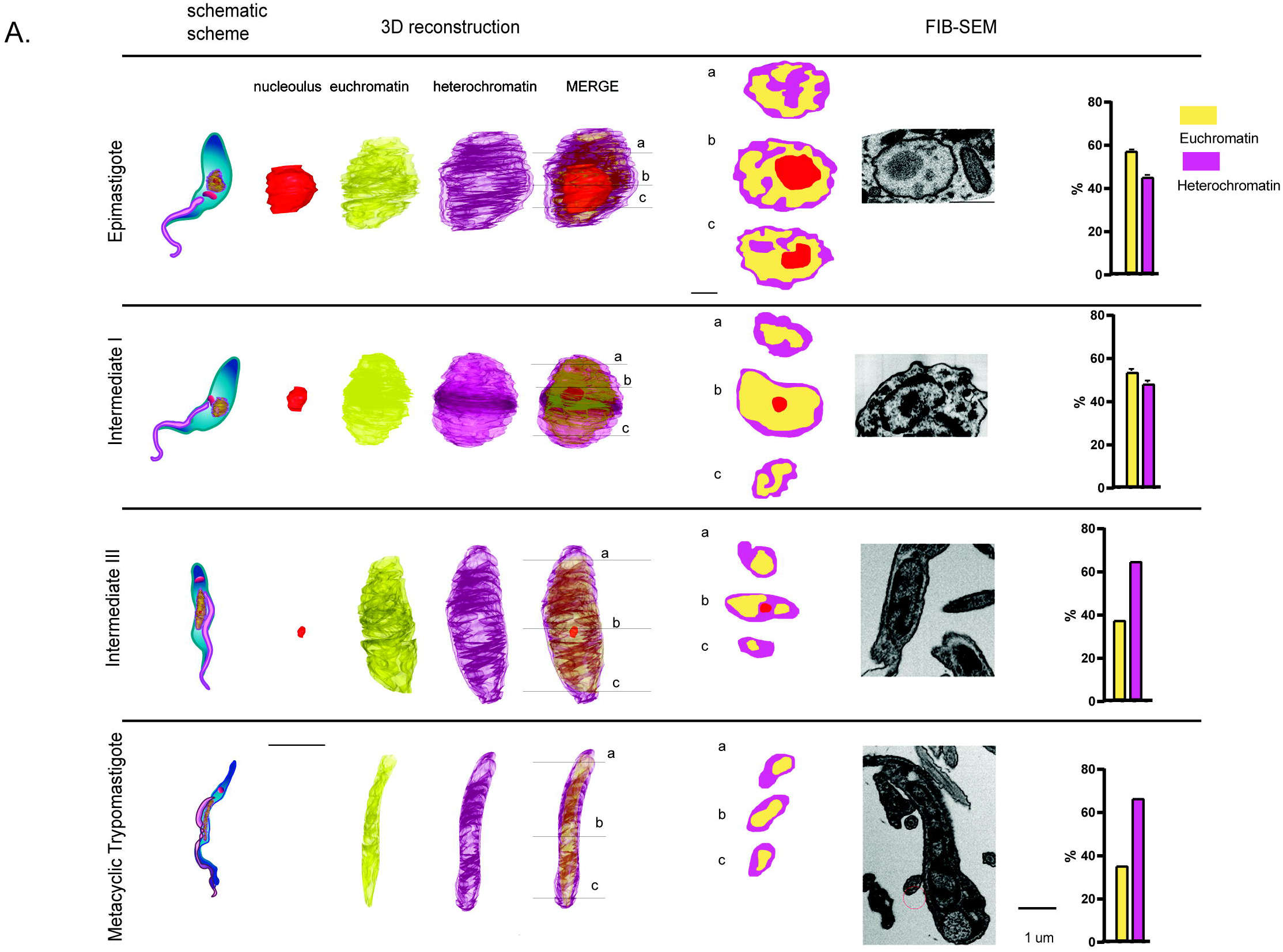
Tridimensional reconstruction of nuclear chromatin regions during *T. cruzi* metacyclogenesis. 3D reconstruction of different developmental stages, from epimastigote to MT, where three different FIB-SEM slices were used to show chromatin state and distribution. High and low electron-dense chromatin regions, which correspond to heterochromatin and euchromatin, respectively, were quantified from TEM slices. Note chromatin remodeling as the differentiation process advances. The euchromatin region (yellow) decreases, whereas the heterochromatin area (purple) increases and occupies mainly the nucleus periphery.

### Genome-wide analysis of open chromatin regions in epimastigotes and MTs

Tridimensional reconstruction of chromatin areas along epimastigote to MT differentiation indicated a significant reduction in euchromatin regions (**Figure 3**), which are considered open chromatin areas. To gain more insights into these changes, we performed a comparison of FAIRE-seq data from epimastigote and MT forms. Visual inspection of normalized RPGC levels along the *T. cruzi* genome confirms a greater abundance in open chromatin and few clear enrichments in epimastigote forms than MTs (**Figure 4A**).

**Figure 4.**
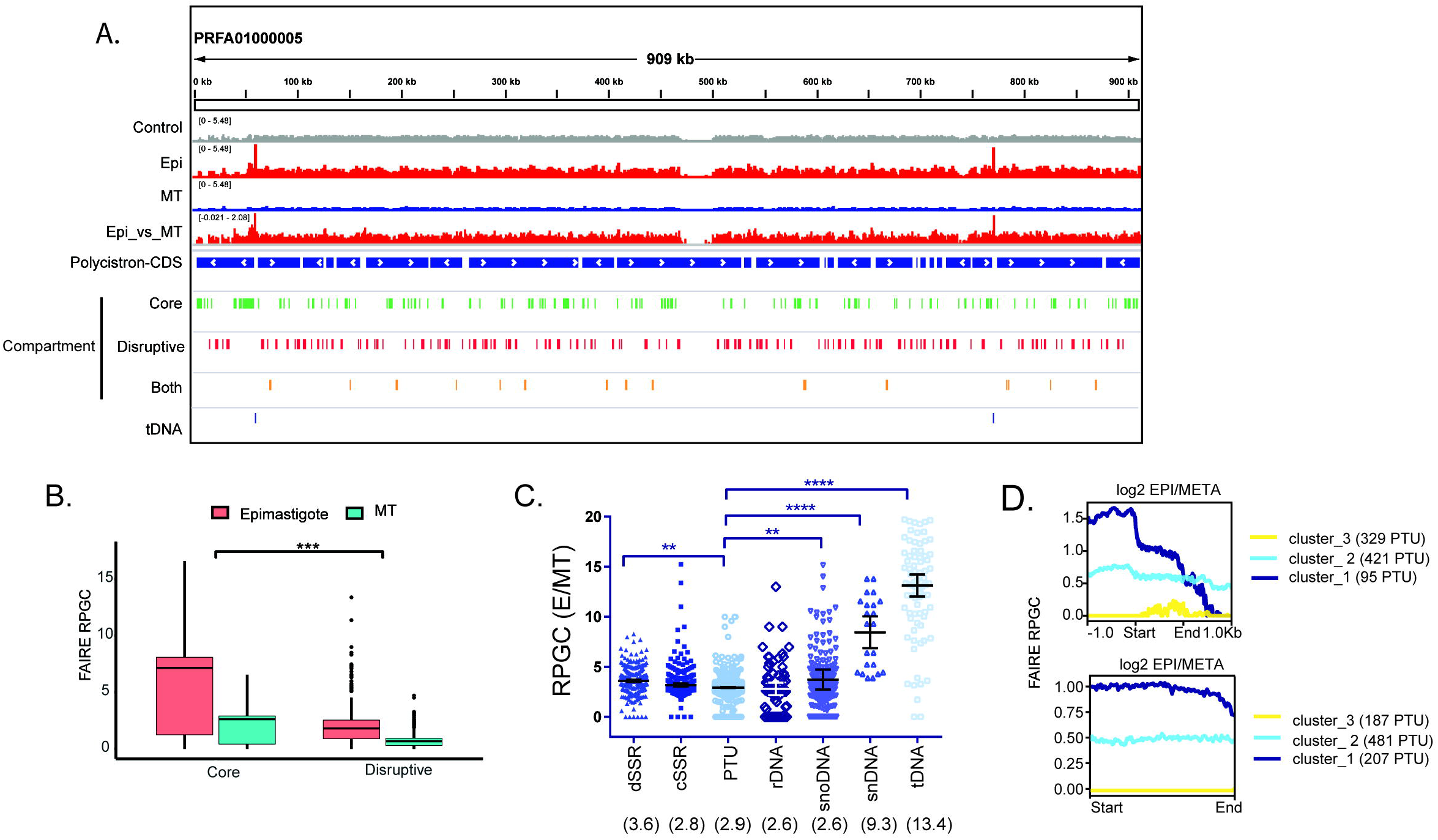
Open chromatin profile changes in life forms. **A.** IGV snapshot of contig PRFA01000005 showing FAIRE-seq profile distribution in epimastigote (red), MT (blue), and control samples (gray). PTUs are shown in blue tracks with the transcription direction indicated by arrows. Genes from disruptive (red), core (green), and those from both (orange) compartments are depicted. tDNAs are shown in dark blue. **B.** Comparison of core and disruptive compartments using RPGC counts. *** Wilcoxon- Mann-Whitney test with p-value = 0.001. **C.** Scatter plots of E/MT RPGC levels on each indicated feature. Median values are written between parentheses. One-way ANOVA with Dunnett’s correction. **D.** Hierarchical cluster analysis of the distribution of the RPGC log2 ratio (E/MTs) in PTUs considering (top) or not (below) 1 kb upstream or downstream. The number of members in each cluster is depicted next to the graph. The first base of the feature is represented by Start, while the last base is End.

As mentioned above, the disruptive compartment is enriched in virulence factors that are mainly expressed in infective forms (23). To address whether the disruptive compartment would be enriched in open chromatin in MTs relative to epimastigotes, RPGC-normalized FAIRE-seq data were compared between life forms, obtaining log2 ratios. Median RPGC values from the disruptive and core compartments were lower in MTs (**Figure 4B, Table S2-3**). Differences between compartments within life forms are very similar: the core compartment has 3.9 times more open chromatin than the disruptive compartment in both epimastigotes and MTs. In the MNase data, core and disruptive compartments also presented different RPGC levels, with the former being significantly higher (**Figure S6**). In general, virulence factors (TS, GP63, MASP, and mucins) and core compartments have 2.6 and 2.8 times (fold change, median values) less open chromatin in MTs than in epimastigotes (**Figure S6**), which agrees with global changes in other genomic features (see **Figure 4C, Table S4**). The open chromatin landscape of virulence factor genes is similar in both life forms, with a slight difference at the upstream coding region of mucins (**Figure S7H**). Taken together, these results indicate that virulence factors indeed have a different pattern of open chromatin when compared to other CDSs (as we discussed below) that seemed to be maintained between life forms, which reinforces major posttranscriptional control of these genes.

Previously, we found that dSSRs from TCTs were enriched in nucleosomes compared to epimastigotes (25). Then, we speculated whether dSSRs from epimastigotes would be more abundant in open chromatin. Indeed, dSSRs are approximately 3.6 more enriched in open chromatin at epimastigotes (**Figure 4C**) (**Figure S8**), which corroborates a lower nucleosome occupancy in replicative forms compared to nonreplicative forms.

Upon hierarchical cluster analysis, the enrichment of open chromatin at dSSRs can be further visualized in a PTU context (**Figure 4 D** **– top, Table S5**). Most notably, Cluster 1 showed enrichment at dSSRs followed by a decreasing signal level toward the cSSRs. A detailed inspection of Cluster 1 elements revealed that approximately 10% of their PTUs are located downstream of tDNA loci. Cluster 2 represents PTUs whose dSSRs have similar levels of openness compared to their adjacent PTUs. Furthermore, interestingly, Cluster 3 encompasses PTUs whose levels of open chromatin are equal between E and MTs (log2 E/MT=0). Taken together, the obtained data indicate that different PTUs have distinct levels of openness at their transcription initiation regions. Clusters 2 and 3 were enriched in genes from the dispersed compartment (**Figure S9**). Functional annotation analysis of the genes from these clusters with or without multigenic family members is shown in **Figure S10**.

Considering only the PTU region, epimastigotes showed a significantly increased signal (2.9 times) compared to MTs (**Figure 4C**). The hierarchical cluster analysis reflected PTUs with different levels of open chromatin among life forms and, importantly, near-flat enrichment along the PTU region (**Figure 4D** **bottom, Table S6**). In accordance with hierarchical clustering of PTUs with SSRs (**Figure 4D** **top**), Clusters 2 and 3 were also enriched in the multigenic family (**Figure S9B**). Cluster 1 was enriched in terms of “cellular nitrogen compound metabolism”, Cluster 2 “endonuclease activity”, “calcium ion binding”, and Cluster 3 “catalytic activity, acting on RNA” (**Figure S11**). The latter is enriched in the SLAC gene, a retrotransposon located close to mini-exon spliced leader genes.

### FAIRE-seq data correlate with steady-state gene expression levels in both life forms

Given the different open chromatin profiles observed among PTUs (**Figure 4D**), we investigated whether a similar pattern occurs in distinct CDSs from different life stages. Thus, CDSs were hierarchically clustered into three groups based on their log2 RPGC level (**Figure S12**). An increase in open chromatin was found mainly at genomic regions encompassing 5’ and 3’ UTRs at Clusters 1 and 2. Similar to the results shown above, Cluster 1, which has the highest log2 ratio, was enriched in genes from the core compartment, whereas Clusters 2 and 3 were enriched in genes from the disruptive compartment and repetitive genes (**Figure S12B**).

Then, we explored whether regions enriched in open chromatin would have transcripts expressed at higher levels. CDSs were first classified as high, medium or low expressed in each life form based on their TPM counts obtained from a transcriptomic study published elsewhere (19) (**Figure S13**). RPGC normalized counts were retrieved for each CDS according to their respective expression class and each life form (**Figure 5**). Surprisingly, a positive correlation of open chromatin with steady-state transcription levels was found. For epimastigotes, significant differences in FAIRE enrichment were observed when comparing all expression classes. In contrast, for MTs, significance was observed mainly when compared to the low expressed genes (high versus low and medium versus low).

**Figure 5.**
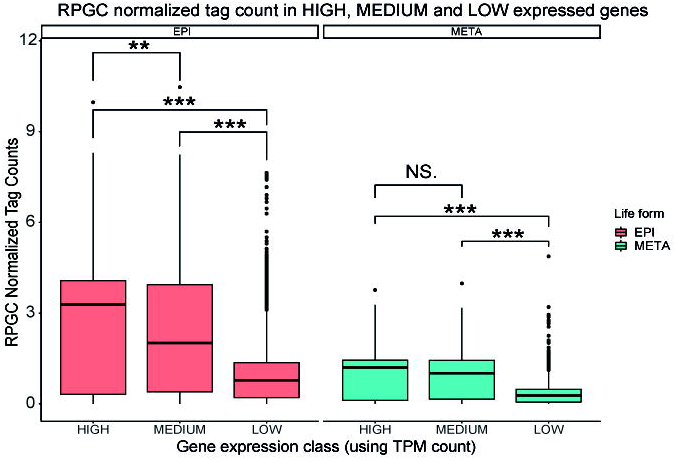
FAIRE-seq data correlate to steady-state transcription levels in both life forms. RPGC normalized tag counts for each gene are mapped to expression classes for epimastigotes (red) and metacyclic trypomastigotes (blue). Statistical significance tests were performed with the Wilcoxon-Mann–Whitney test (for p values: *** = 0.001; ** = 0.01; N. S = Not Significant).

### Open chromatin is developmentally regulated at tDNA loci, likely reflecting translation regulation

FAIRE-seq analysis highlighted a global decrease in the levels of open chromatin in MTs compared to epimastigotes. However, we wondered whether the open chromatin profile would give us more clues about the differences in gene expression and phenotype found between life forms, especially those related to their differentiation program. Comparing all genomic features, we detected striking differences in open chromatin enrichment at the regions uncovered by the small nuclear RNAs (snRNAs) and tRNA genes. Of note, this enrichment was significantly higher (fold changes of 13.4 and 9.3 for tDNAs and snDNA, respectively) in epimastigotes than in MTs (**Figure 4C and 6A-B**).

**Figure 6.**
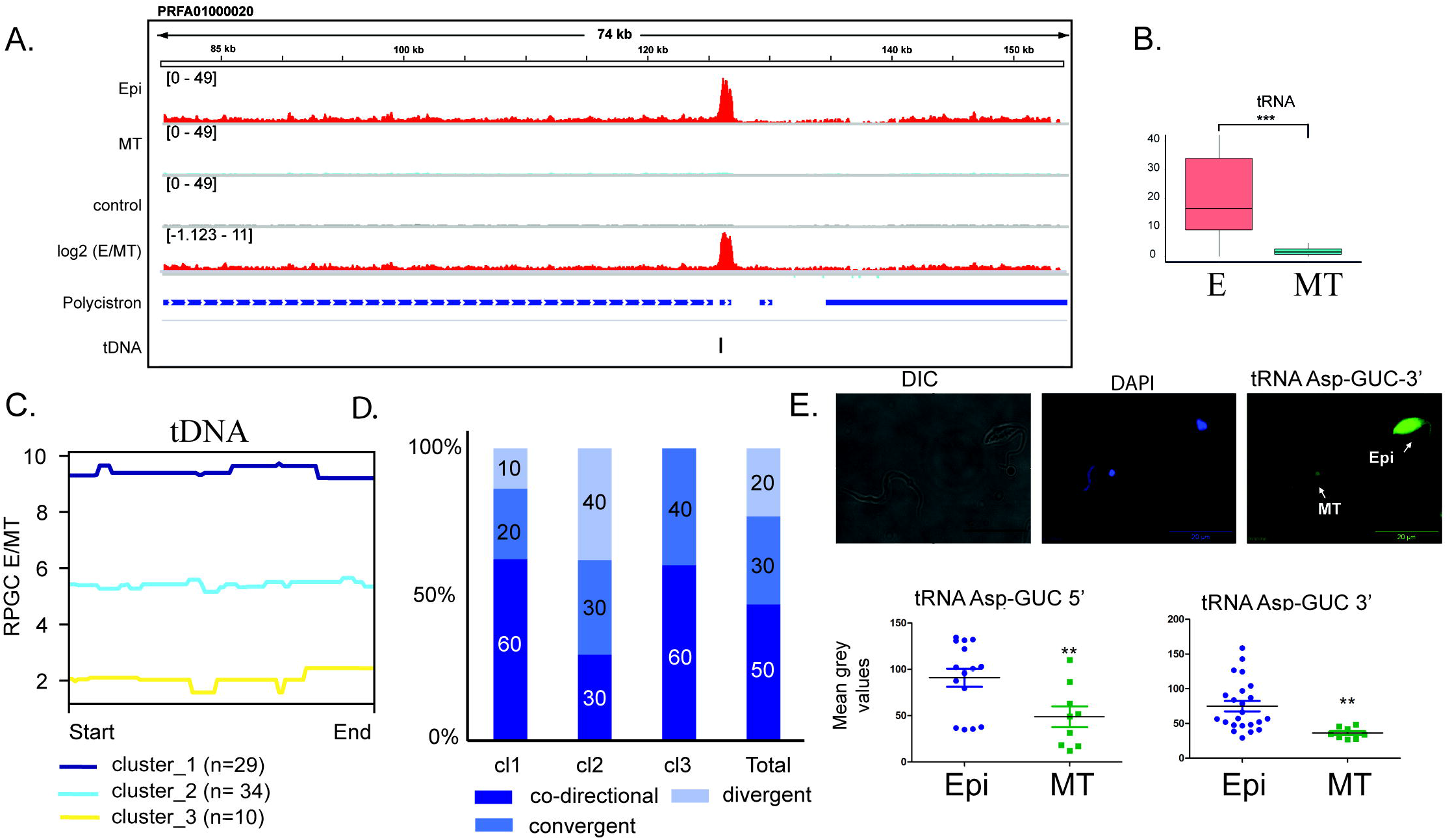
FAIRE enrichment at tDNA loci. **A.** IGV snapshot of a representative tDNA cluster (black box) showing FAIRE enrichment in epimastigote (red) tracks over metacyclic trypomastigotes (blue). **B.** Boxplot of RPGC normalized tag counts in tDNA features (*** Wilcoxon-Mann-Whitney test with p-value = 0.001). **C.** Hierarchical cluster analysis of FAIRE-seq data at tDNA loci (reciprocal ratio). The number of tDNAs in each cluster is depicted below. **D.** Distribution of tDNAs from each cluster of **C** according to their location relative to the adjacent PTU transcription direction. **E.** RNA-FISH analysis of Asp-GUC tRNAs in epimastigotes and MTs. Bottom, quantification of tRNA-GUC 5’ and tRNA-Asp-GUC 3’ by FISH in epimastigote (blue) and MT (green) forms. Mean gray values represent intensity per area. Unpaired T-test: ** (p-value > 0.05).

Hierarchical cluster analysis of the E/MT ratio revealed that the majority of tDNAs had at least six times more open chromatin in epimastigotes than in MTs (**Figure 6 C, Figure S14, Table S7**). Clusters were not related to the tRNA isoaceptor type or class (**Figure S16A**); however, their distribution reflects their location regarding the transcription direction of the adjacent PTU (**Figure S15B**). For example, Clusters 1 and 2 were enriched in tDNAs that were mainly located between codirectional (60% and 30%, respectively) and divergent (10% and 40%, respectively) PTUs (**Figure 6D**). This distribution suggests that chromatin alterations at tDNAs may affect transcription initiation at adjacent PTUs.

Finally, we addressed whether the increase in open chromatin within tDNAs would be associated with tRNA expression levels. Evaluation of tRNA expression by PCR is not a straightforward approach once this molecule is highly modified. Then, we evaluated tRNA expression in life forms using an RNA FISH approach. Previously, (29) showed that compared to MTs, epimastigotes have much higher amounts of tRNAs. To confirm this finding, epimastigotes and MTs were probed simultaneously using one of the four probes against 5’ and 3’ Asp and Glu tRNAs. **Figures 4E** **and S16** indicate that tRNAs are much lower in MTs, indicating that tRNAs are approximately 50% less expressed in nonreplicative forms.

## Discussion

The differentiation of *T. cruzi* into nonreplicative life forms is followed by a global decrease in RNA transcription, which is mainly observed in fully differentiated forms, such as TCTs and MTs (17, 18). Chromatin is the stage where gene expression regulation occurs, and trypanosomes offer an exciting scenario to evaluate its changes once their protein-coding genes are devoid of transcription regulation in a gene-specific manner (6). Here, we showed that the differentiation of epimastigotes to MTs is followed by a progressive increase in the percentage of heterochromatin area, which becomes restricted to the nuclear periphery (Figure 3). Using a conformation chromosome capture approach, euchromatic and heterochromatic regions were further associated with structural genome compartments A and B, which are related to transcriptionally active and inactive regions, respectively (30, 31). This work indicates that the differentiation program that drives morphological (such as repositioning of kinetoplast and flagellum, nuclear elongation) and metabolic changes in trypanosomes, is also followed by intense remodeling of chromatin domains.

To further explore the genomic context of open chromatin, we used the FAIRE approach, a methodology previously known to enrich for active genomic regions (3,5,32). Then, we infer that dSSRs are enriched in open chromatin relative to PTUs, corroborating the presence of fewer nucleosomes at dSSRs than PTUs (25). Interestingly, the *T. cruzi* genome has PTUs whose dSSRs have different levels of open chromatin. Whether it is associated with differential deposition of the RNA Pol II machinery, histone variants, and histone PTMs and, more importantly, whether it has any consequences on the PTU transcription levels needs to be further investigated. Global nascent transcriptomic analysis comparing transcription along and within PTUs is absent in trypanosomes but can clarify this issue. In addition, hierarchical cluster analysis of PTUs without SSRs indicates different levels of open chromatin between PTUs but with a clear flat pattern along the entire PTU. The latter indicates a uniform open chromatin status along PTUs, possibly reflecting the polycistronic transcription nature of trypanosomes.

We envisaged that the presence of two genomic compartments in *T. cruzi*, proposed by Berná *et al.*, may induce the formation of a complex 3D chromatin structure. Here, by evaluating open chromatin status in a genome-wide fashion, the core and disruptive genomic compartments of *T. cruzi* exhibited great differences in the enrichment of open chromatin, with the former highly enriched in open chromatin compared to the latter. The disruptive compartment is composed of virulence factors that have a very repetitive nature. Therefore, we cannot rule out that mapping and genome assembly bias influence these data, although removing multimapping reads did not affect the results. Regardless, because virulence factors are mainly expressed in infective forms, it was expected that these chromatin regions would be more open in MTs if a gene-specific transcriptional regulation scenario existed. However, it was determined that both the disruptive and core compartments were lower in MTs. These findings confirm that posttranscriptional regulation of virulence factors indeed plays a key role in their expression, as previously indicated (33, 34).

Moreover, the multigenic families MASP, GP63, mucin, and TS exhibited distinct open chromatin landscapes compared to the other nonmultifamily members. The reason for this may be related to the greater GC content of the disruptive compartment than the core compartment (22), because the G plus C content is intrinsically correlated with nucleosome occupancy (35). Although an expected opposite correlation among FAIRE and MNase-seq data was observed in the majority of evaluated genomic regions, clearly it was possible to note that some regions do not contain nucleosomes but are depleted in active chromatin and vice versa, indicating the presence/absence of other chromatin players. An important variable is the fact that kinetoplastids contain a thymine modification called base J that is mainly located in telomeric regions. In *T. cruzi*, base J was also found flanking PTUs and within subtelomeric regions, which are rich in virulent factors (located at the disrupted compartment). Interestingly, the presence of base J at these genes was shown to be developmentally regulated (36, 37). Thus, it remains to be investigated whether the presence of base J at the disruptive compartment may interfere with chromatin openness and/or if other protein complexes may be in these regions, thereby affecting chromatin architecture.

Strikingly, the open chromatin state based on FAIRE-seq read enrichment is associated with steady-state transcription levels in *T. cruzi*. These findings are remarkable once trypanosome gene expression relies mainly on posttranscriptional events. However, this is not unprecedented. Previously, it was determined that the nucleosome landscape reflects steady-state gene expression mainly regarding the depth of the nucleosome-depleted region around 5’UTR (7,25,38). A proposed explanation is the association of cotranscriptional splicing resulting in putative interference with the final levels of transcripts. It can be also hypothesized that the machinery associated with RNA processing (trans-splicing and polyadenylation) and some RNA binding proteins would closely associate with chromatin, thereby influencing local chromatin structure. In fact, *T. cruzi* chromatin proteomics reveals the presence of at least 13 mRNA binding proteins (24), but further experiments should be performed to determine whether these proteins indeed interact with chromatin and if they may play a role in chromatin architecture.

Of note, the open chromatin landscape seems to be conserved between life forms, except for a global decrease at their levels on MTs. This agrees with a global decline in transcription levels along with parasite differentiation (18). One of the main differences in FAIRE enrichment between life forms was found at loci that code for snRNAs and tRNAs. snRNAs are uridine-rich sequences that compose the spliceosome and are transcribed by RNA polymerase III (MARCHETTI et al., 1998). tDNAs are approximately 78 bp long and contain intrinsic promoters, known as Box A and Box B, which are the binding sites for the transcription factor complex named TFIIIC and are responsible for recruiting the TFIIIB complex to enable the assembly of RNA polymerase III at the TSS region of these genes (41, 42). Due to the higher expression of tDNAs on epimastigotes, we believe that the RNA Pol III complex would be enriched in tDNA loci, likely hampering the deposition of nucleosomes and generating an open chromatin status. In fact, epimastigotes indeed have a lower enrichment of nucleosomes at tDNAs (compared to TCTs). Nevertheless, in TCT life forms, tDNA loci are depleted of nucleosomes compared to their surrounding regions (25).

Of note, the absence of transcription machinery at tDNA loci can be the cause or consequence of the closed chromatin status. However, the implications of the developmental regulation of this status can be hypothesized. The mentioned changes may influence the expression of the nearby PTUs, which, in turn, would be preferentially affected along with differentiation. In this sense, it is interesting that tDNA hierarchical cluster analysis pointed out a distinct distribution of tDNA openness based on their location related to adjacent PTUs (convergent, divergent, or codirectional). We hypothesize that the loss of open chromatin status at tDNAs located within a codirectional PTU may affect transcription mainly at the downstream PTU. This is supported by the findings that indicate that tDNA loci and their associated transcription factors, mainly TFIIIC, may function as insulators. The latter has a critical role in chromatin domain organization, working on separating active and silenced domains (43).

Developmentally regulated chromatin changes at tDNA loci are especially intriguing considering that translation control has a key role in trypanosomatid differentiation. During the differentiation of epimastigotes to MTs, a global downregulation of a large proportion of the translatome occurs (19,21,44). Although the underlying mechanism is not fully understood, phosphorylation of eiF2a increases upon nutritional stress and attenuates protein synthesis during *T. cruzi* differentiation (21). Here, we confirmed that tRNAs are less expressed in MTs than in epimastigotes, correlating with the diminished amount of open chromatin in the former than in the latter. Thus, chromatin changes at tDNA loci may be another layer of translation regulation during development. In fact, many studies indicate that tRNA levels are indeed a limiting step in translation (45), blocking it when presented below a given threshold.

## Conclusion

Overall, our genome-wide approach recapitulates unique aspects of trypanosomatid gene regulation such as polycistronic transcription and lack of RNA pol II transcription regulation. However, it also indicated that different dSSRs may have different levels of open chromatin, suggesting PTU-specific regulation. In addition, the status of open chromatin correlates with the presence of conserved and disruptive genome compartments, which further correlates with the expression of virulence factors. Finally, the obtained data indicated developmental chromatin regulation at tDNA loci, which may orchestrate an essential step in posttranscriptional regulation control necessary for parasite differentiation and survival.

## Methods

### Parasites

*T. cruzi* (Dm28c strain) epimastigotes were cultured at 28 °C in LIT medium supplemented with 10% fetal bovine serum (FBS - Vitrocell), glucose 0.4%, hemin 0.1 µM, and penicillin-G 59 mg/L as previously described (46). Metacyclic trypomastigote (MT) parasites were obtained by the metacyclogenesis protocol described at (47). Briefly, epimastigotes in the exponential growth phase were diluted at 1×10^6^ parasites/mL and cultivated for five days to reach the stationary phase. Then, parasites were resuspended at 5×10^8^ parasites/mL in triatomine artificial urine medium (TAU, 190 mM NaCl, 17 mM KCl, 2 mM CaCl_2_, 2 mM MgCl_2_, 8 mM phosphate buffer pH 6.0) for 2 hours at 28 °C. Next, parasites were diluted to 5×10^6^ per mL in TAU medium supplemented with 10 mM L-proline, 50 mM L-glutamate, 2 mM L-aspartate, and 10 mM glucose and maintained for 96 hours at 28 °C. MTs were obtained from the supernatant for 72 hours and purified by DEAE columns.

### Nuclear tridimensional reconstruction

Image stacks previously obtained by focused ion beam scanning electron microscopy (FIB-SEM) (15) were automatically aligned using the StackReg plugin in the Fiji/ImageJ software package (http://fiji.sc/wiki/index.php/Fiji). Aligned images were saved as image sequence files and then imported into the 3D reconstruction program Free-D 1.15 (48). Heterochromatin, euchromatin, and nucleolus were accomplished by manual classification and tracing the boundary contours of ultrastructures of interest. Visualization of the 3D shape of the cell nucleolus (red), heterochromatin (purple), and euchromatin (yellow) was achieved by representing the traces of these objects using Free-D 1.15. Nuclear 3D reconstructions were performed in triplicate using 70 to 130 slices for each cell.

### FAIRE sample preparation

The FAIRE samples were prepared as described in (3) with some modifications. A total of 2 x 10^7^ parasites (in biological duplicates) were fixed for 5 min with formaldehyde and directly added to the medium to a final concentration of 1%. Formaldehyde was quenched with 125 mM glycine, followed by extensive washing with PBS. Pellets were resuspended in 1 mL of cold lysis buffer TELT (50 mM Tris-HCl pH 8.0, 62.5 mM EDTA, 2.5 M LiCl, 4% Triton X-100) and sonicated with six cycles of 30 s with an interval of 1 min each (Output 4 and Duty 30 parameters) in a Tommy Ultrasonic Disruptor UD-201 apparatus. To prepare the input DNA control, 100 μL of each sample was centrifuged at 15,000 g for 5 min at 4 °C. The supernatants were incubated with 1 μL of RNaseA DNase-free for 30 min at 37 °C followed by treatment with 1 μL of proteinase K for 1 h at 55 °C. The cross-links were reversed by overnight incubation at 65 °C. DNA was extracted with phenol:chloroform:isoamyl alcohol solution and precipitated with 3 M sodium acetate (pH 5.2) and 95% ethanol. To prepare FAIRE DNA, the remaining cell lysate was centrifuged at 15,000 for 5 min at 4 °C, and DNA was extracted as described above; however, RNaseA treatment, protein removal by proteinase K and cross-linking reversal were performed after DNA purification. The obtained DNA was quantified using a NanoDrop spectrophotometer (Thermo Fisher). A total of 500 ng of input DNA control was run on a 1% agarose gel to verify the sonication efficiency. DNA fragments were deep sequenced using an Illumina NextSeq 500 (2 x 75 bp) at the Polyomics Facility (Glasgow University).

### Bioinformatic analyses

#### Sequence read quality check, mapping, and normalization

The methods applied to analyze the FAIRE-seq data are shown in Figure S1 C and described below. Raw sequencing files were quality checked with FASTQC (49). A quality filter was not necessary (see Figure S1 B). Reads were then mapped against the *Trypanosoma cruzi* Dm28c genome (downloaded from https://tritrypdb.org/ - DB46, Dm28c 2018 version) using the GATK pipeline (50), which employs BWA (51) as the alignment algorithm. The GATK pipeline was executed until reaching the read deduplication step. To avoid ambiguous read mapping, the alignment files were filtered using a MAPQ (MAPing Quality) score of 10 using SAMtools (52). MAPQ filtered alignment files were processed with multiBamSummary (binSize = 50) from deepTools 3.3.0 (53) to generate the scaling factors for normalization and the files necessary to perform PCA and Spearman correlation analysis (using plotPCA and plotCorrelation functions, respectively). The data were normalized using the RPGC (Reads Per Genome Coverage) method, present in bamCoverage function, applying the following scaling factors for each dataset: EpiR1: 2.3087, EpiR2: 4.0037, MetaR1: 0.5875, MetaR2: 0.6076, ControlR1: 0.6763, ControlR2: 0.6551. Replicates were merged into one bigwig file using the bigWigCompare function, calculating the mean signal for each replicate. The comparisons Epi vs. Control and Epi vs. Meta were performed with the bigWigCompare function using the parameters skipZeroOverZero, pseudocount 1 1, and operation log2. The datasets generated and/or analyzed during the current study are available in the NCBI BioProject repository under the accession code PRJNA763084 and BioSample codes SAMN21423261, SAMN21423262, SAMN21423263, SAMN21423264, SAMN21423265 and SAMN21423266.

#### Extragenomic feature annotation

Polycistronic regions (same strand CDS cluster region), dSSR and cSSR were annotated using the GFF file from *T. cruzi* Dm28c (downloaded from https://tritrypdb.org/ - DB46, Dm28c 2018 version) and a custom Python script (available at: https://github.com/alexranieri/annotatePolycistron). Briefly, the script parses the GFF file, and a polycistron is considered for annotation if two or more CDSs of the same strand direction are found consecutively. The ID of each feature is incorporated in the polycistron annotation content, along with the number of features found. Polycistrons with only one feature were also annotated. In a second analysis, the script processes the SSRs, and dSSR is considered when a region between the start of a negative-strand polycistron followed by the start of a positive-strand polycistron is found. cSSR is considered when a region between the end of a positive-strand polycistron followed by the end of a negative-strand polycistron is found. If none of those conditions are met, i.e., a positive-strand CDS polycistron is followed by another type of feature. The genomic region between them is annotated as an Intergenic Polycistron Region.

5’ and 3’ UTR annotations were obtained with UTRme (54) using the spliced-leader sequence (AACTAACGCTATTATTGATACAGTTTCTGTACTATATTG) from (55) and transcriptomic data from Smircich et al., 2015. tRNAscan-SE (56), with default parameters for eukaryotes, was used to identify anticodons and amino acids from tRNA sequences obtained from hierarchical cluster analysis.

#### Transcriptomic analysis

RNA-seq reads were preprocessed according to Smircich et al., 2015, with an additional step of sequencing adapter removal using cutadapt (57). Then, the processed reads were mapped against the *T. cruzi* Dm28c (V46) genome using STAR (58). For each CDS in the genome, transcripts per million (TPM) values were obtained from transcriptomic data using TPMCalculator (59). RPGC normalized tag counts (in log2) were retrieved using annotatePeaks function from HOMER (60), passing the wig files, CDS coordinates in bed file and the parameters: -size ‘given’, - raw or -noadj (to avoid new normalization). Genes were classified into low, medium, or high expressed by ordering TPM values from highest to lowest and then divided into three groups containing the same elements. CDS IDs were correlated to obtain both TPM and RPGC tag count values. Statistical significance tests between value counts (TPM and RPGC) of expression classes were performed with the Wilcoxon-Mann-Whitney test.

#### Identification of open chromatin landscapes and visualization

To avoid plotting regions from small sequences, only features from the largest contigs were selected. Thus, the first 188 sequences of the genome were analyzed, which comprised 85% of the total number of bases. Genomic coordinates of each feature analyzed were obtained from the final GFF annotation (after using the annotatePolycistron Python script and UTRme) as bed files. Landscape profiles and heatmaps were plotted using the computeMatrix (with scale-regions and skipZeros options) and plotHeatmap (with hierarchical clustering) functions from deepTools. The final GFF annotation, bed files, and RPGC normalized data (as wig files) were used as input in IGV (Integrative Genomics Viewer) (61) for additional inspection and visualization.

#### Additional analysis

Gene Ontology (GO) analysis was performed by passing the feature IDs to TriTrypDB and then using the GO Enrichment tool. The obtained results were exported to REVIGO (62) for visualization. Repeat regions were obtained from http://bioinformatica.fcien.edu.uy/cruzi/browser/downloads.php.

#### Superposition of FAIRE and MNAse-seq data

MNase-seq data were retrieved and processed according to our previous study (25). The aligned bam files were RPGC normalized using bamCoverage from deepTools, applying the following scaling factors for each sample: SAMN16241276: 1.0625, SAMN16241277: 0.9521, SAMN16241279: 0.9223, SAMN16241280: 1.0565, SAMN16241281: 1.0104, and SAMN16241282: 0.9837.

### tRNA Fluorescence *in situ* Hybridization (FISH)

Parasites (10^7^) in the exponential growth phase were fixed with 4% paraformaldehyde in PBS for 20 min at room temperature. Pellets were resuspended in NH_4_Cl under agitation for 10 min, washed with PBS, and adhered to slides with an adhesive frame previously treated with polylysine. Then, parasites were permeabilized with 0.2% Triton X-100 in PBS for 5 min, blocked and prehydrated with 2% BSA, 5X Denhardt, 4X SSC and 35% formamide for 2 h. A total of 1 ng per mL of fluorescent oligonucleotide was added to the hybridization solution: 1 mM EDTA, 1% BSA, 0.5 M NaHPO_4_, 7% SDS, and heated at 75 °C for 5 min. For hybridization, a wet chamber was prepared with 50% formamide in 2X SSC, and the slides inside were incubated at 37 °C overnight. The slides were washed twice with 4X SSC, four times with 50% formamide and 2X SSC, three times with 2X SSC, three times with 1X SSC, and three times with 0.5X SSC. The slides were mounted with Vectashield plus DAPI solution (Vectors). Images were captured by an Olympus BX51 microscope, and the intensity of fluorescence was quantified by ImageJ with mean gray values.

## Declarations

Ethics approval and consent to participate

Not applicable

## Consent for publication

Not applicable

## Availability of data and materials

The datasets generated and/or analyzed during the current study are available in the NCBI BioProject repository under the accession code PRJNA763084 and BioSample codes SAMN21423261, SAMN21423262, SAMN21423263, SAMN21423264, SAMN21423265 and SAMN21423266.

## Competing interests

The authors declare that they have no competing interests.

## Funding

This work was supported by fellowships from FAPESP and by grants (#18/15553-9, #18/14432-3, #13/07467-1, #19/19690-3, # 2019/19834-5, # 20/02708-4) from the Sao Paulo Research Foundation (FAPESP), by the Serrapilheira Institute (grants Serra- 1709-16865 and Serra-1709-16607), and Fundação de Amparo à Pesquisa do Estado do Rio de Janeiro (FAPERJ). ARJL and HMCS were supported by fellowships from FAPESP; SBP was supported by a fellowship from the Serrapilheira Institute. MCE has a fellowship from the National Council for Scientific and Technological Development (CNPq). The sponsors have no role in the design of the study and collection, analysis, and interpretation of data or in writing the manuscript.

## Authors’ contributions

JPCC, MCE and SB conceived the original idea. SB, JNR, HMCS and LL performed the wet experiments. ARL processed and analyzed the computational data. JNR, HMCS, HGS, ARJ, LL and JPCC analyzed the data. FH, PAG, MCMM, CM, MCE and AMS helped supervise the project. JPCC supervised the project. JPCC and ARL wrote the manuscript. All authors provided critical feedback and helped shape the research, analysis and manuscript.

## Supporting information

Legends for Supplemental Figures

## Acknowledgments

We thank Karin Cruz and Ivan Novaski Avino for their technical assistance. We thank Cibele Masotti for reading this manuscript and providing critical comments, Dr. Jose Patané for initial statistical analysis, and Dr. David Silva and Daniel Ohara for bioinformatics support.

## Legends for Supplemental Figures

**Figure S1. A**. Data summary for sequenced samples (EPI – epimastigote; META – metacyclic trypomastigote) and control for each replicate and read pairs. **B.** FastQC mean quality scores of sequenced samples. **C.** Scheme of the pipeline used to explore the FAIRE-seq data **D.** Number of mapped reads against the *Trypanosoma cruzi* Dm28c genome for each dataset (biological replicates). Blue bars represent total dataset reads; orange bars indicate the mapped reads; gray bars show the remaining mapped reads with MAPQ scores above 10. The percentage is relative to the total reads of each dataset (blue bars).

**Figure S2.** Spearman correlation (**A**) and PCA (**B**) of read counts for each dataset (biological replicates) from epimastigote (EPI), metacyclic trypomastigote (META) life forms and control generated with deepTools.

**Figure S3.** Representative IGV snapshot of FAIRE-seq data mapped against the *T. cruzi* Dm28c genome (contig PRFA01000010). The data were normalized using RPGC (see Materials and Methods in main text). Epimastigote biological replicates (EPI-1 and EPI-2) are shown in red, and metacyclic trypomastigote biological replicates (META-1 and META-2) are shown in blue. Datasets not filtered with MAPQ score 10 are labeled as “WithoutQ10”. At the genomic features, repeat regions are represented in yellow. The dashed rectangle highlights a long repeat region and its low coverage upon MAPQ10 removal.

**Figure S4.** IGV snapshots of two contigs comparing the raw mapped FAIRE read profile to the removal of multimapped reads (Q10 - arrows) in epimastigotes and metacyclic trypomastigotes. Core and disruptive genomic compartments are shown as red and green tracks, respectively.

**Figure S5.** Superposition of FAIRE-seq (red) and MNase-seq (blue) in epimastigotes. The y-axis represents RPGC normalized tag counts for FAIRE-seq on the right side and MNase-seq on the left side. The profile of other CDSs transcribed by RNA Pol II is represented in **A**, and multigenic family gene profiles are represented in **B-G**.

**Figure S6.** Comparison of core and disruptive compartments of MNase-seq data using RPGC counts in epimastigotes (red) and TCTs (blue). The median values for epimastigores were 2.04 and 1.09 for the core and disruptive compartments, respectively; for TCT, the median values were 2.13 and 1.14. *** Wilcoxon-Mann-Whitney test with p-value = 0.001.

**Figure S7. A-F**. RPGC tag counts at multigenic family genes and other CDSs transcribed by RNA Pol II (**G**) for epimastigotes (red) and metacyclic trypomastigotes (blue). H. Sumary plots for virulence factors in epimastigotes and MTs. *** Wilcoxon- Mann-Whitney test with p-value = 0.001.

**Figure S8.** Landscape profile of cSSR (blue) and dSSR (green) showing median values in RPGC log2 ratio of epimastigotes/MTs (left). Start and End represent the first and last bases of each feature, respectively. Boxplots showing RPCG counts for dSSR (upper) and cSSR (lower) for epimastigotes (red) and MTs (blue). *** Wilcoxon- Mann–Whitney test with p value = 0.001

**Figure S9.** The percentage of bases of multifamily genes for each polycistron according to clustering is shown in Figure 4D. In **A**, clusterization was performed considering 1 kb upstream and downstream from the polycistron, while in **B**, clusterization considered only the PTU. Statistical analysis was performed using the Wilcoxon-Mann-Whitney test (*** p-value = 0.001; * p-value = 0.05).

**Figure S10.** Functional annotation analysis (GO – Molecular Function and Biological Process) of genes from hierarchical cluster analysis of PTUs with SSRs after removal of multifamily genes (Figure 4D)

**Figure S11.** Functional annotation analysis (GO – Molecular Function and Biological Process) of genes from hierarchical cluster analysis of PTUs without SSRs after removal of multifamily genes (Figure 4D)

**Figure S12.** Hierarchical cluster and function analysis of CDS. **A.** Landscape profiles of all CDSs (upper) and clustering based on the RPGC log2 ratio of epimastigotes/MTs (lower). **B.** Functional GO annotation of biological processes using RPGC log2 ratio clustering

**Figure S13.** Genes classified as high, medium and low expressed based on TPM values for epimastigote (red) and metacyclic trypomastigote (blue) life forms. *** Wilcoxon- Mann-Whitney test with p-value = 0.001

**Figure S14.** IGV snapshots of FAIRE-seq enrichment in epimastigotes (duplicates – red) compared to MTs (duplicates – blue) in three different loci. tDNAs are highlighted in yellow rectangles.

**Figure S15. (A)** Number of tRNA genes in each cluster. The tRNA genes grouped in each cluster after hierarchical clustering, as shown in Figure 6C, were counted and plotted in column graphics according to R group classification of the amino acids. (B) Representative scheme of tRNA gene localization at divergent, convergent and codirectional PTUs. The arrows indicate the sense of transcription in each PTU.

Figure S16. Quantification of tRNA-FISH UUC in epimastigote (blue) and MT (green) forms. Mean gray values represent intensity per area. Unpaired T-test: ** (p-value > 0.05), and *** (p-value < 0.0001).

## Supplemental videos

**Video S1.** 3D reconstruction of euchromatin (yellow), heterochromatin (purple) and nucleolus (red) in epimastigote stage.

Video S2. 3D reconstruction of euchromatin (yellow), heterochromatin (purple) and nucleolus (red) in intermediate 1 stage.

Video S3. 3D reconstruction of euchromatin (yellow), heterochromatin (purple) and nucleolus (red) in intermediate 2 stage.

Video S4. 3D reconstruction of euchromatin (yellow), heterochromatin (purple) and nucleolus (red) in MT stage.

## Supplemental Tables

**Table S1.** Repetitive regions at *T. cruzi* genome.

**Table S2.** Log2 RPGC at core compartment.

**Table S3.** Log2 RPGC at disruptive compartment.

**Table S4.** RPGC at the indicated features in epimastigotes and MTs (Figure 4C).

**Table S5.** Hierarchical clustering output from Figure 4D_up (PTUs with dSSR and cSSR).

**Table S6.** Hierarchical clustering output from Figure 4D_bottom (PTUs- Start and End only).

**Table S7.** Hierarchical clustering output from Figure 6C (tDNAs).

## Notes

### Competing Interest Statement

The authors have declared no competing interest.

