## Supplemental Figures for "Open chromatin analysis in *Trypanosoma cruzi* life forms highlights critical differences in genomic compartments and developmental regulation at tDNA loci"

A.

| Sample Name | % Duplicates | % GC | Millions of Seqs |
|---|---|---|---|
| EPI-1_R1 | 71.2% | 39% | 43.8 |
| EPI-1_R2 | 64.8% | 39% | 43.8 |
| EPI-2_R1 | 72.1% | 38% | 32.8 |
| EPI-2_R2 | 66.2% | 38% | 32.8 |
| META-1_R1 | 57.4% | 42% | 63.6 |
| META-1_R2 | 52.5% | 42% | 63.6 |
| META-2_R1 | 57.0% | 43% | 61.9 |
| META-2_R2 | 51.7% | 43% | 61.9 |
| Control-1_R1 | 37.7% | 46% | 35 |
| Control-1_R2 | 32.8% | 46% | 35 |
| Control-2_R1 | 38% | 46% | 36.2 |
| Control-2_R2 | 33.3% | 46% | 36.2 |

B.

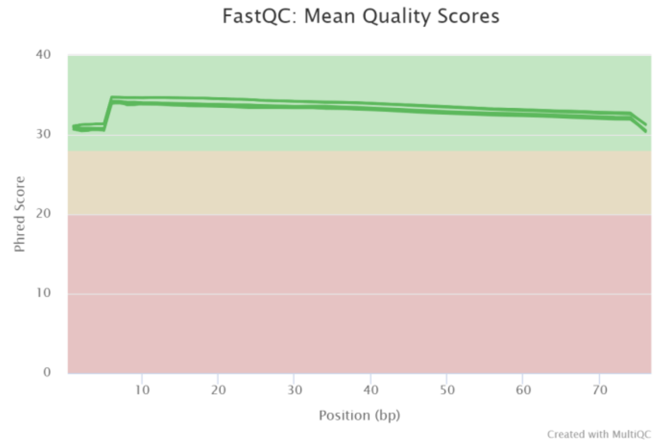


C.

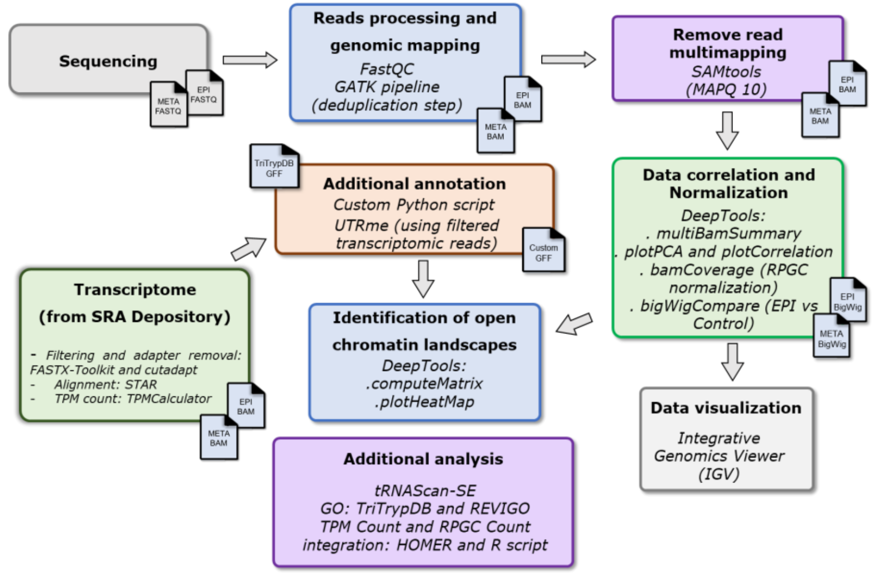


D.

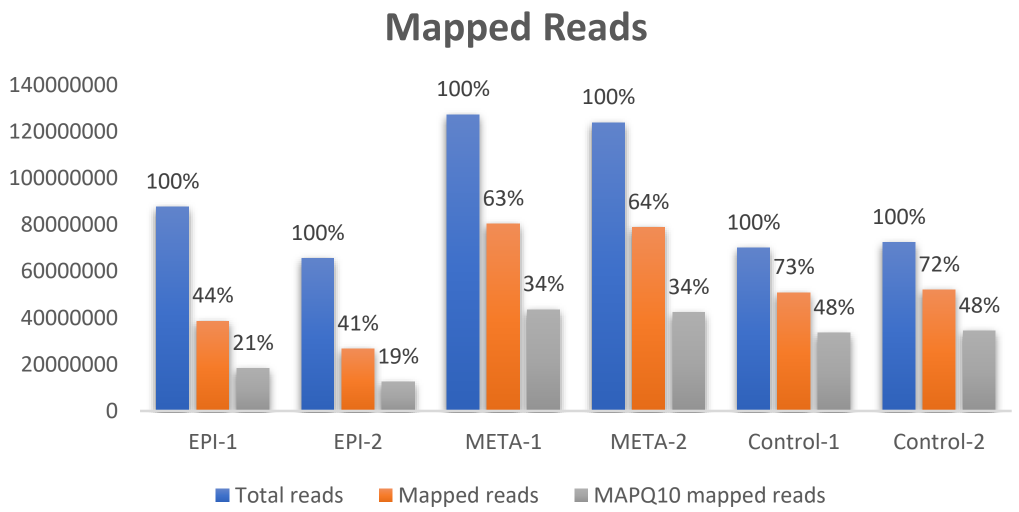


**Figure S1. A**. Data summary for sequenced samples (EPI – epimastigote; META – metacyclic trypomastigote) and control for each replicate and read pairs. **B.** FastQC mean quality scores of sequenced samples. **C.** Scheme of the pipeline used to explore the FAIRE-seq data **D.** Number of mapped reads against the *Trypanosoma cruzi* Dm28c genome for each dataset (biological replicates). Blue bars represent total dataset reads; orange bars indicate the mapped reads; gray bars show the remaining mapped reads with MAPQ scores above 10. The percentage is relative to the total reads of each dataset (blue bars).

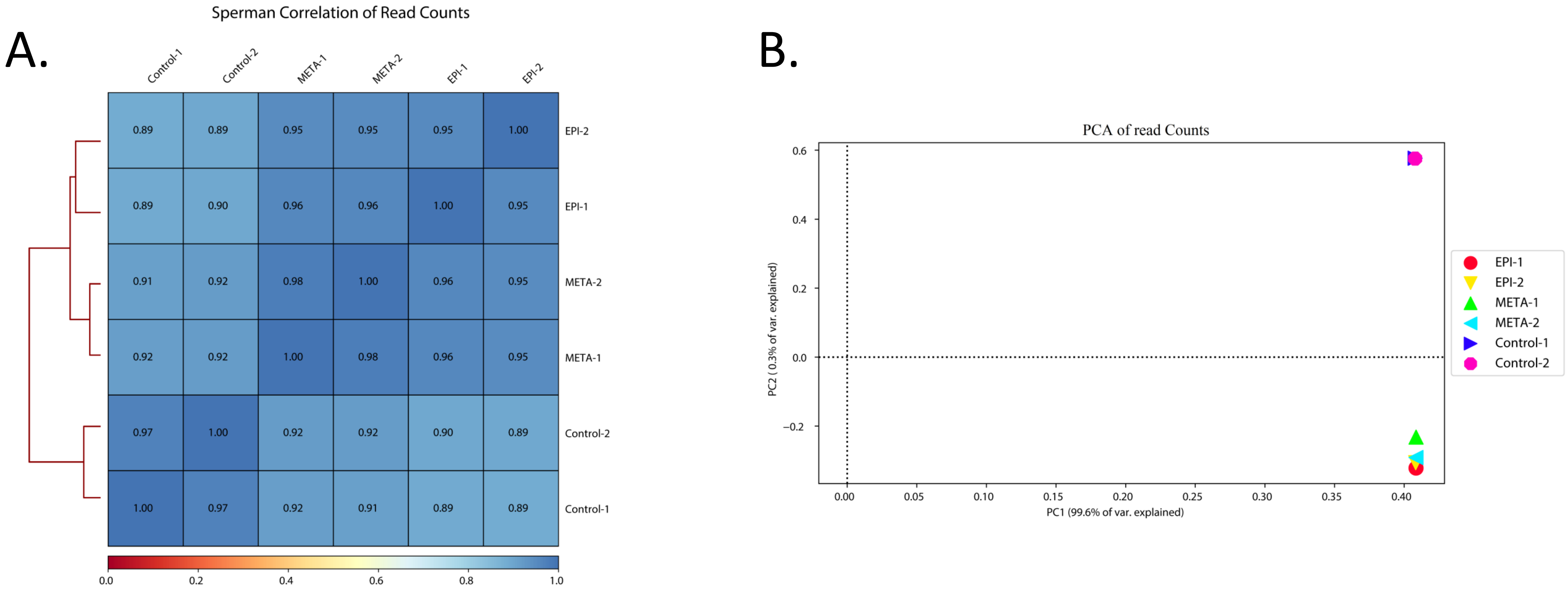


**Figure S2.** Spearman correlation (**A**) and PCA (**B**) of read counts for each dataset (biological replicates) from epimastigote (EPI), metacyclic trypomastigote (META) life forms and control generated with deepTools.

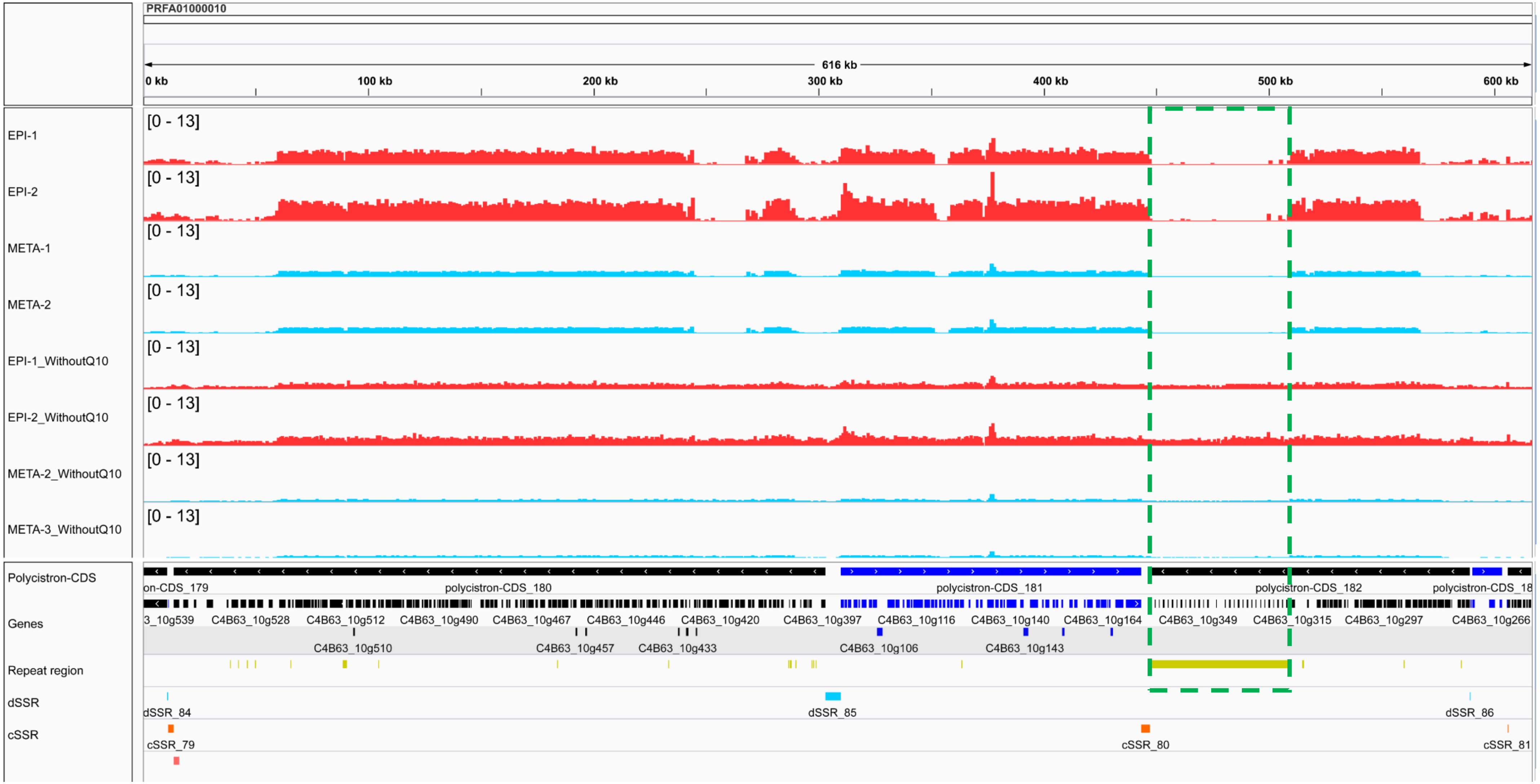


**Figure S3.** Representative IGV snapshot of FAIRE-seq data mapped against the *T. cruzi* Dm28c genome (contig PRFA01000010). The data were normalized using RPGC (see Materials and Methods in main text). Epimastigote biological replicates (EPI-1 and EPI-2) are shown in red, and metacyclic trypomastigote biological replicates (META-1 and META-2) are shown in blue. Datasets not filtered with MAPQ score 10 are labeled as "WithoutQ10". At the genomic features, repeat regions are represented in yellow. The dashed rectangle highlights a long repeat region and its low coverage upon MAPQ10 removal.

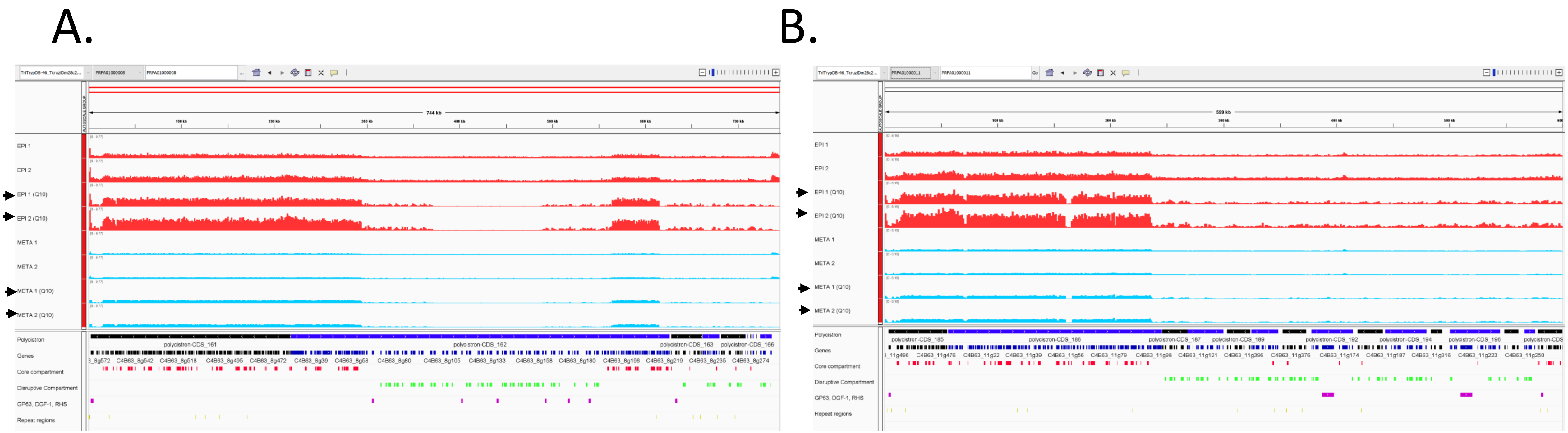


**Figure S4.** IGV snapshots of two contigs comparing the raw mapped FAIRE read profile to the removal of multimapped reads (Q10 - arrows) in epimastigotes and metacyclic trypomastigotes. Core and disruptive genomic compartments are shown as red and green tracks, respectively.

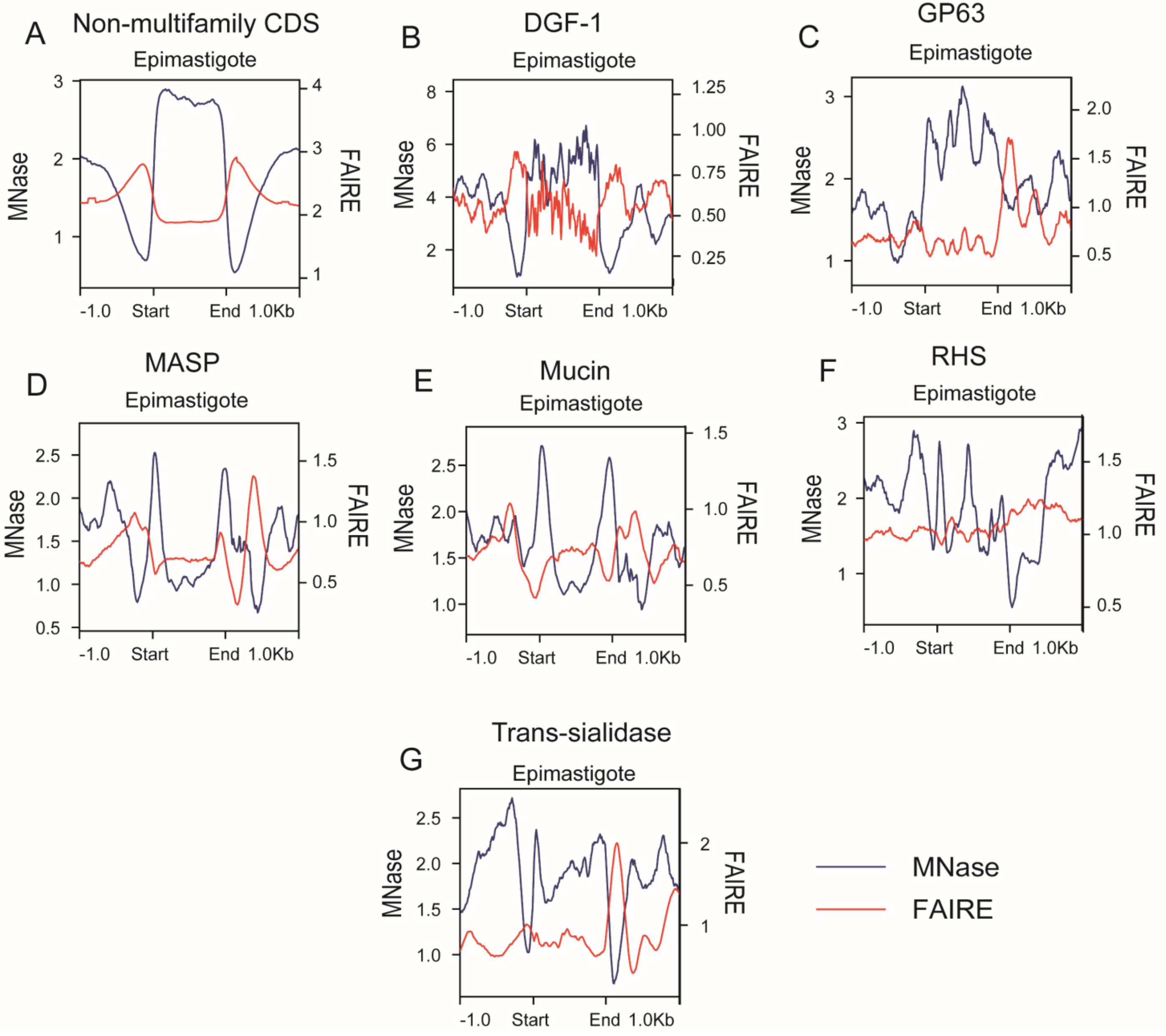


**Figure S5.** Superposition of FAIRE-seq (red) and MNase-seq (blue) in epimastigotes. The y-axis represents RPGC normalized tag counts for FAIRE-seq on the right side and MNase-seq on the left side. The profile of other CDSs transcribed by RNA Pol II is represented in **A**, and multigenic family gene profiles are represented in **B-G**.

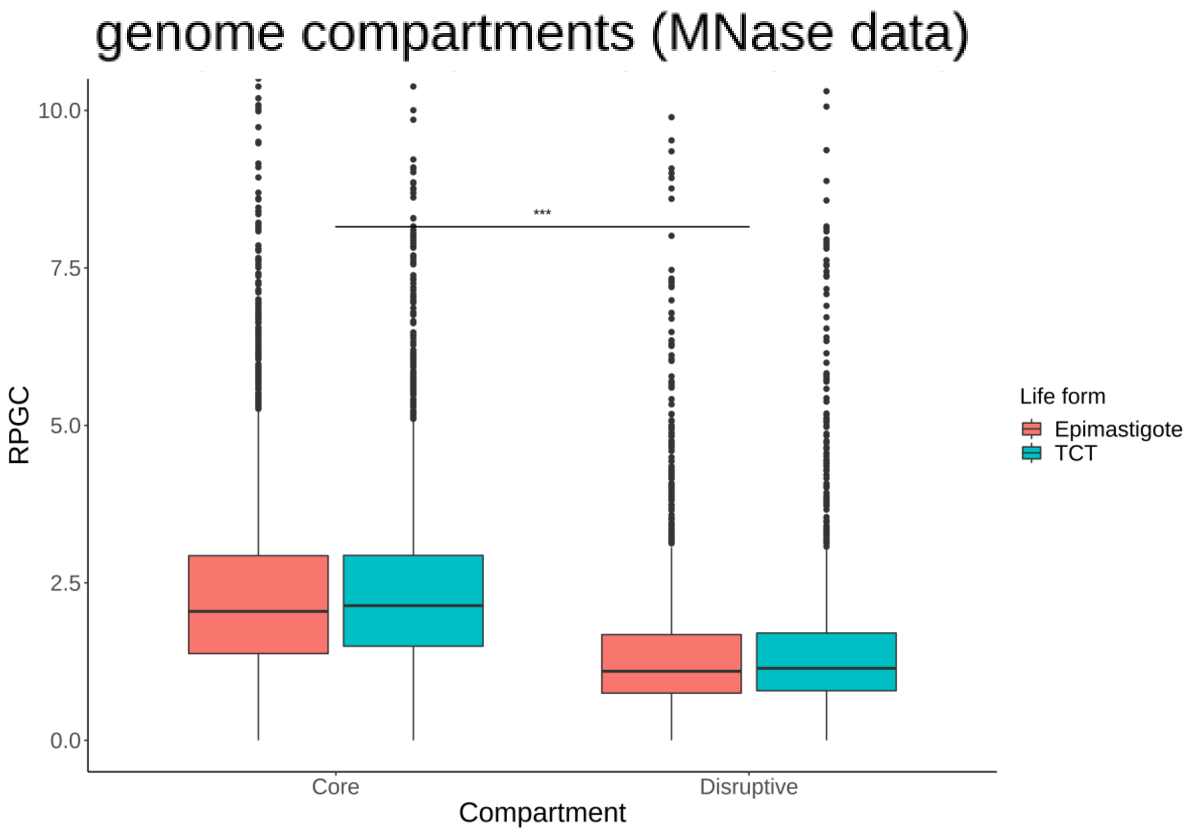


**Figure S6.** Comparison of core and disruptive compartments of MNase-seq data using RPGC counts in epimastigotes (red) and TCTs (blue). The median values for epimastigores were 2.04 and 1.09 for the core and disruptive compartments, respectively; for TCT, the median values were 2.13 and 1.14. *** Wilcoxon-Mann-Whitney test with p-value = 0.001.

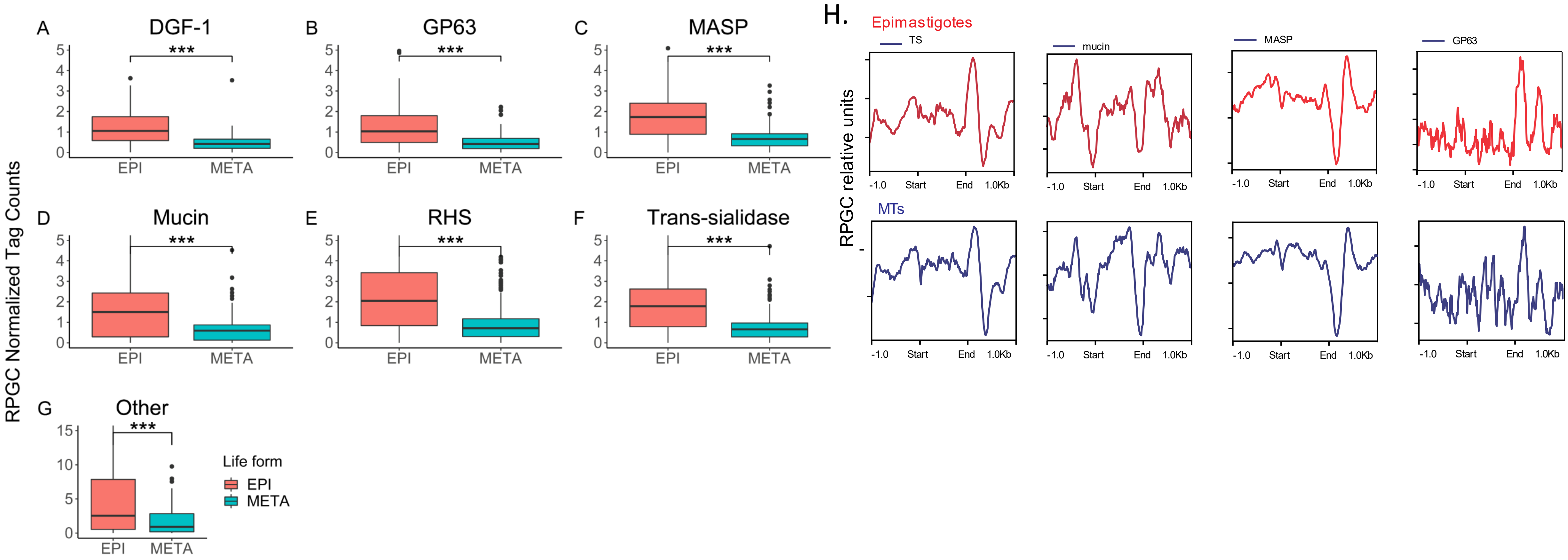


**Figure S7. A-F**. RPGC tag counts at multigenic family genes and other CDSs transcribed by RNA Pol II (**G**) for epimastigotes (red) and metacyclic trypomastigotes (blue). H. Sumary plots for virulence factors in epimastigotes and MTs. *** Wilcoxon-Mann-Whitney test with p-value = 0.001.

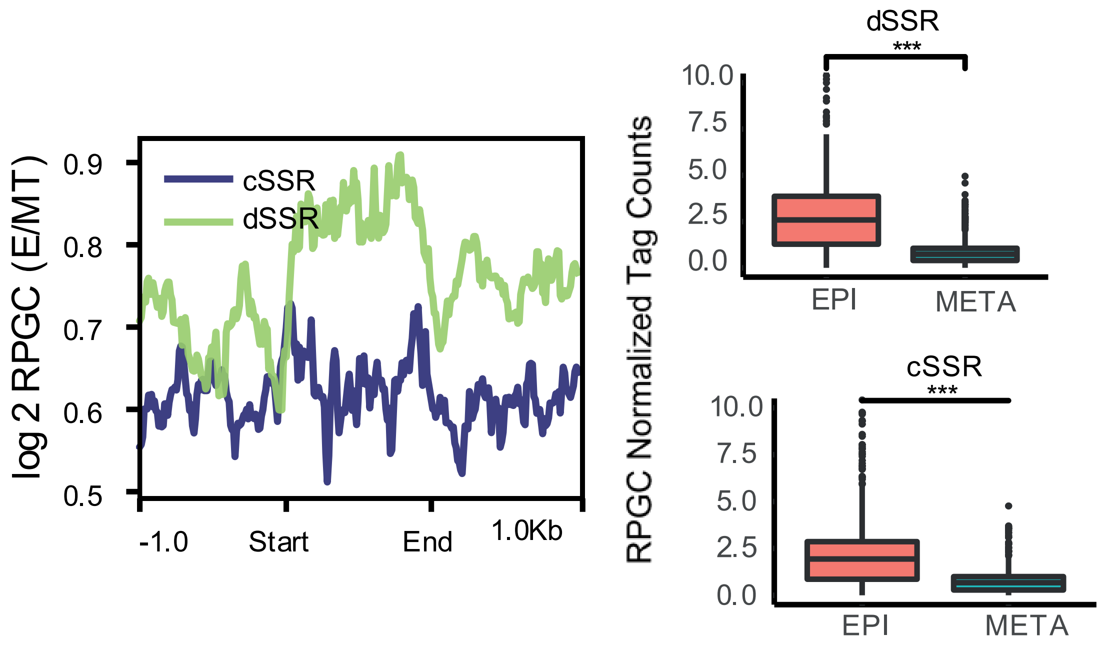


**Figure S8.** Landscape profile of cSSR (blue) and dSSR (green) showing median values in RPGC log2 ratio of epimastigotes/MTs (left). Start and End represent the first and last bases of each feature, respectively. Boxplots showing RPCG counts for dSSR (upper) and cSSR (lower) for epimastigotes (red) and MTs (blue). *** Wilcoxon-Mann–Whitney test with p value = 0.001.

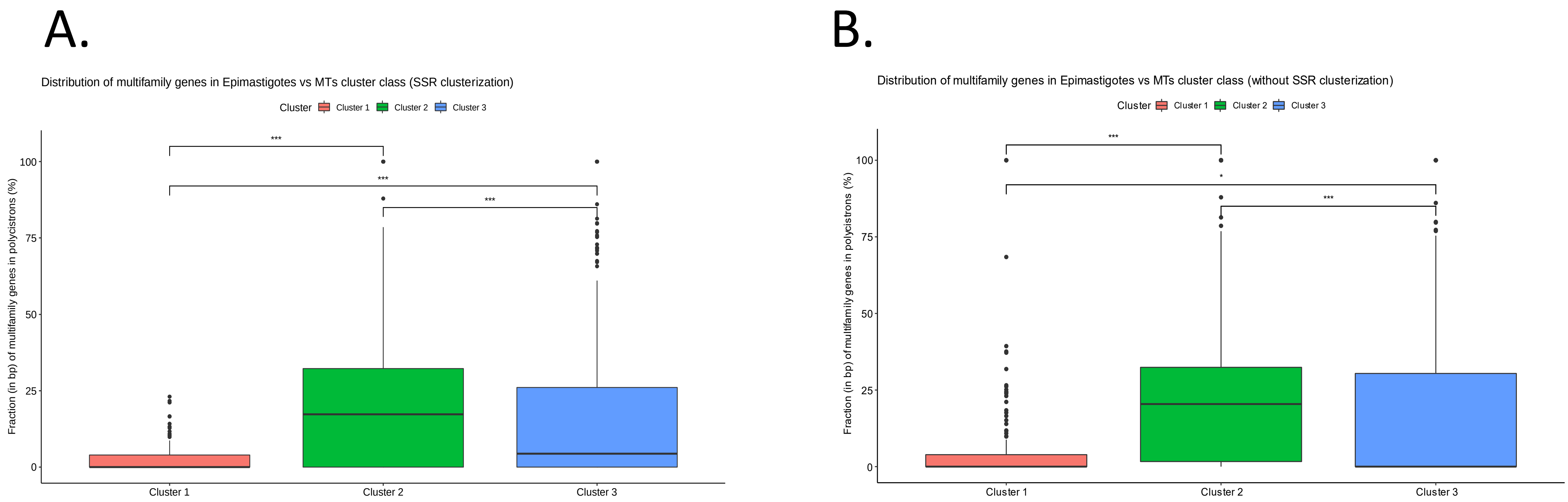


**Figure S9.** The percentage of bases of multifamily genes for each polycistron according to clustering is shown in Figure 4D. In **A**, clusterization was performed considering 1 kb upstream and downstream from the polycistron, while in **B**, clusterization considered only the PTU. Statistical analysis was performed using the Wilcoxon-Mann-Whitney test (*** p-value = 0.001; * p-value = 0.05).

Cluster 1
(4366 genes,
204 MFs)

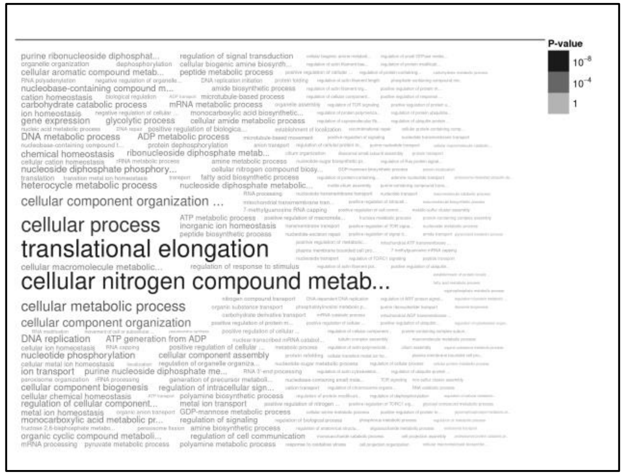


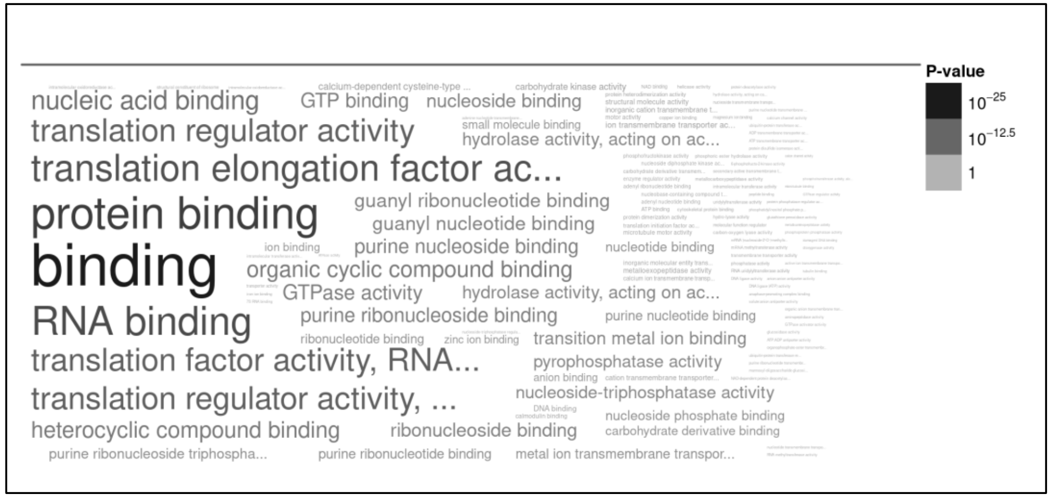


Cluster 2
(6341 genes,
1489 MFs)

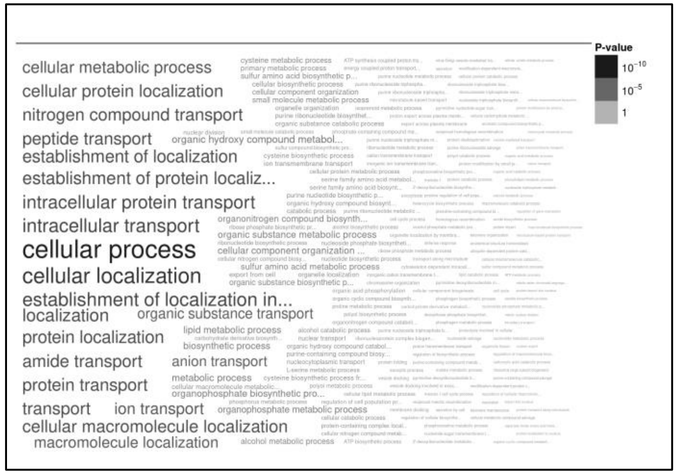


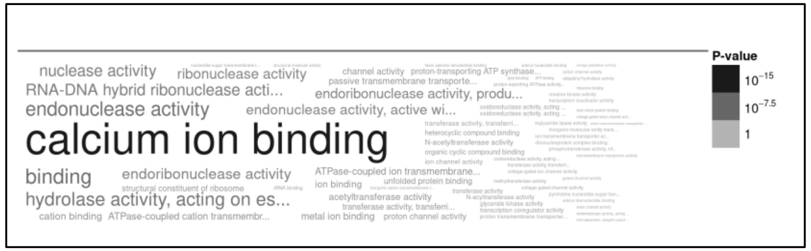


Cluster 3
(4252 genes,
904 MFs)

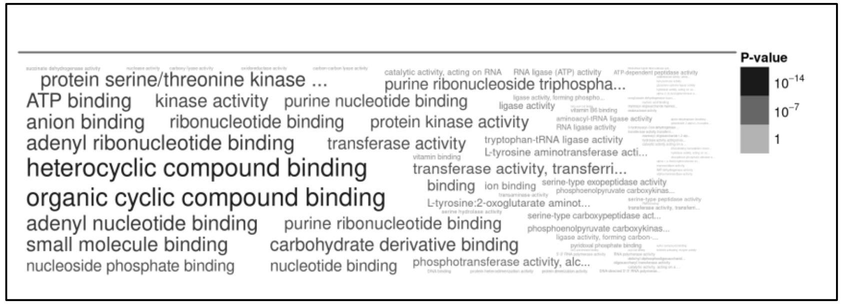


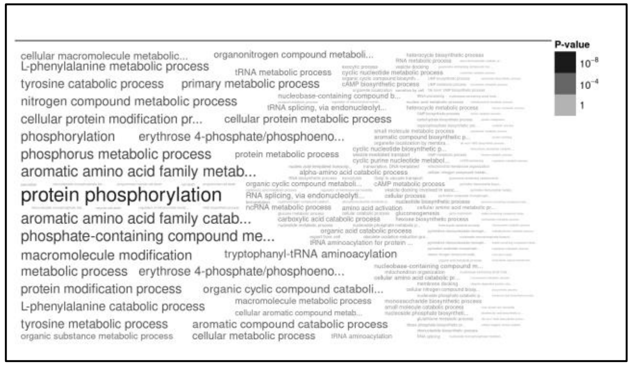


**Figure S10.** Functional annotation analysis (GO – Molecular Function and Biological Process) of genes from hierarchical cluster analysis of PTUs with SSRs after removal of multifamily genes (Figure 4D).

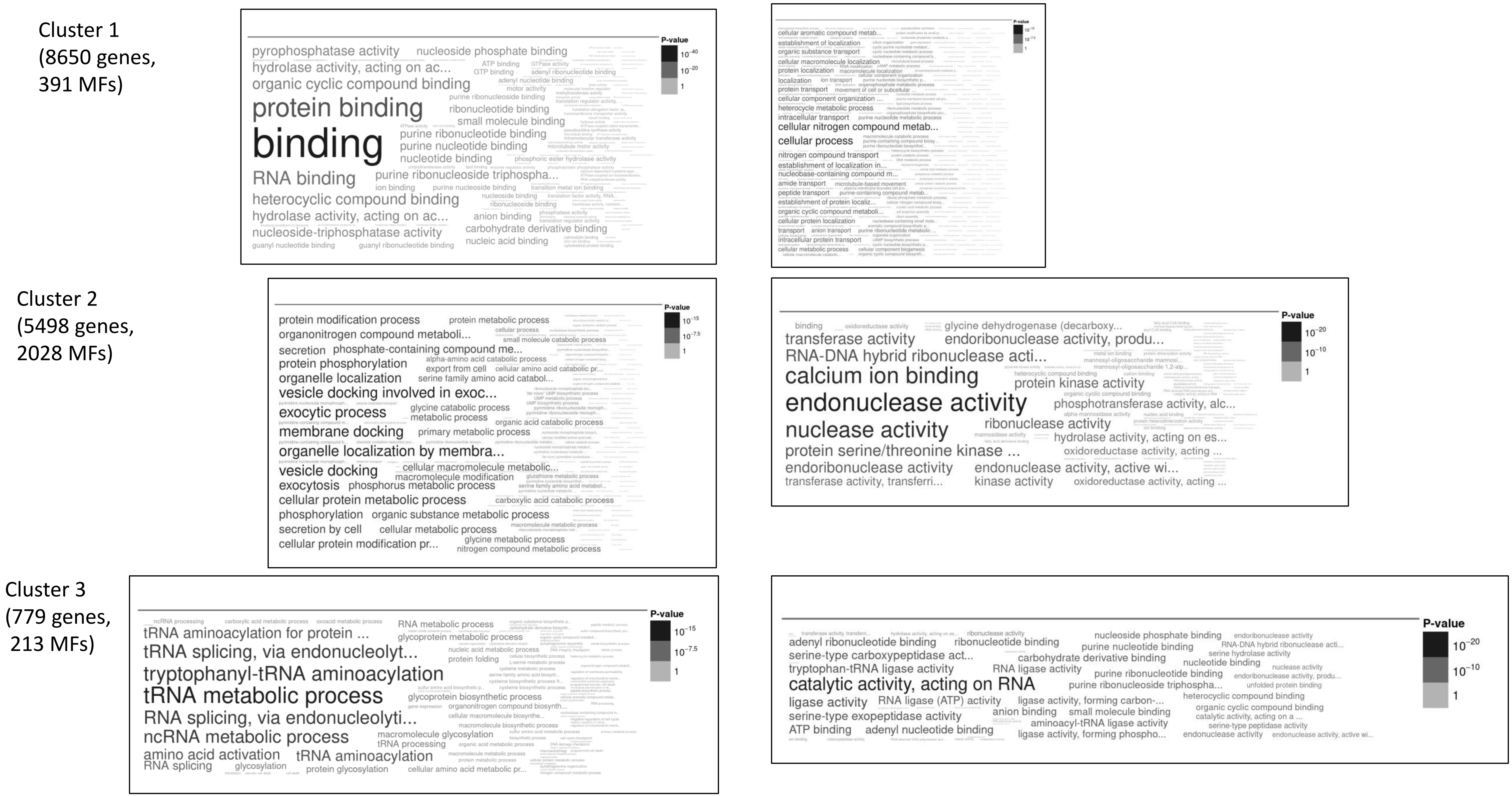


**Figure S11**. Functional annotation analysis (GO – Molecular Function and Biological Process) of genes from hierarchical cluster analysis of PTUs without SSRs after removal of multifamily genes (Figure 4D).

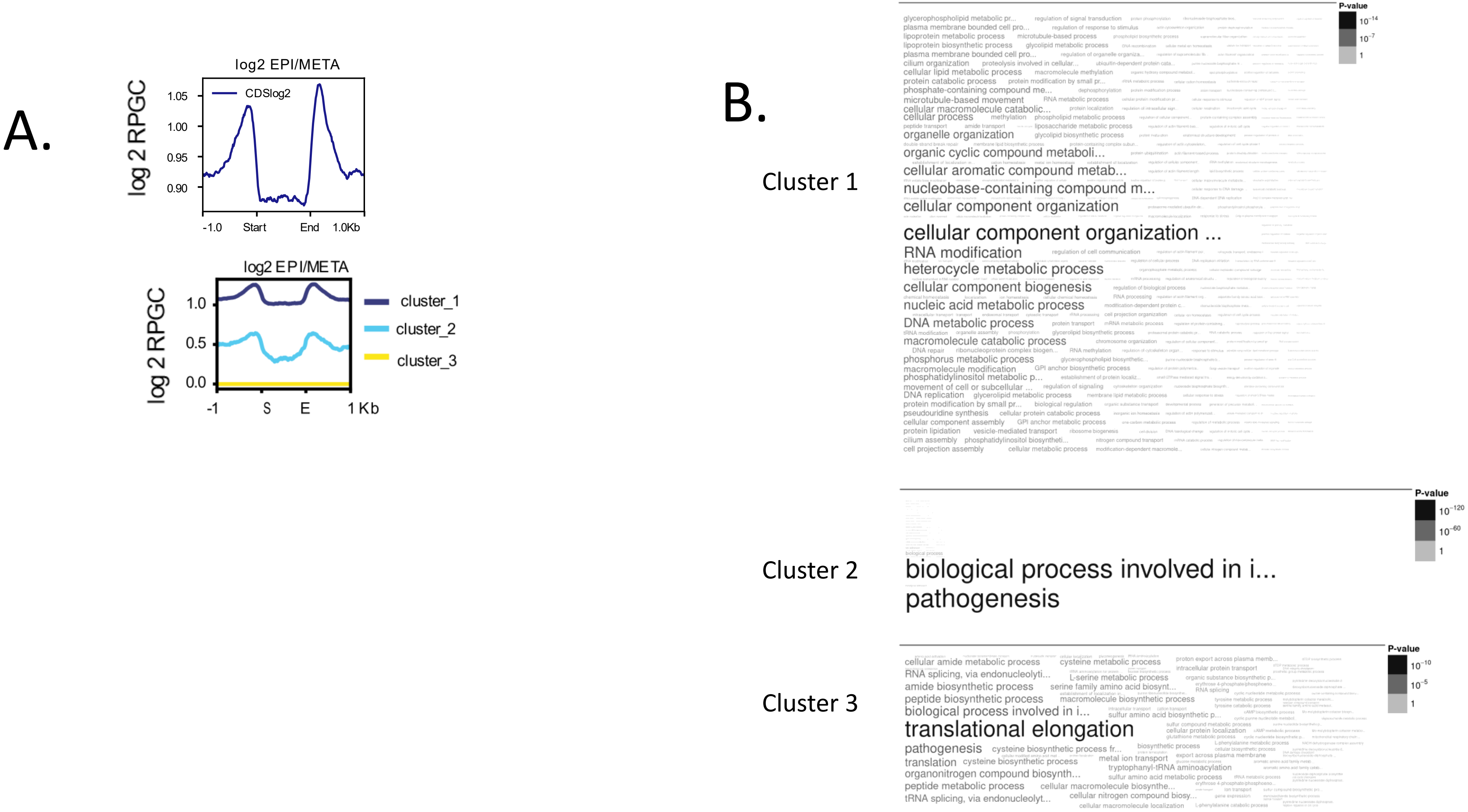


**Figure S12.** Hierarchical cluster and function analysis of CDS. **A.** Landscape profiles of all CDSs (upper) and clustering based on the RPGC log2 ratio of epimastigotes/MTs (lower). **B.** Functional GO annotation of biological processes using RPGC log2 ratio clustering.

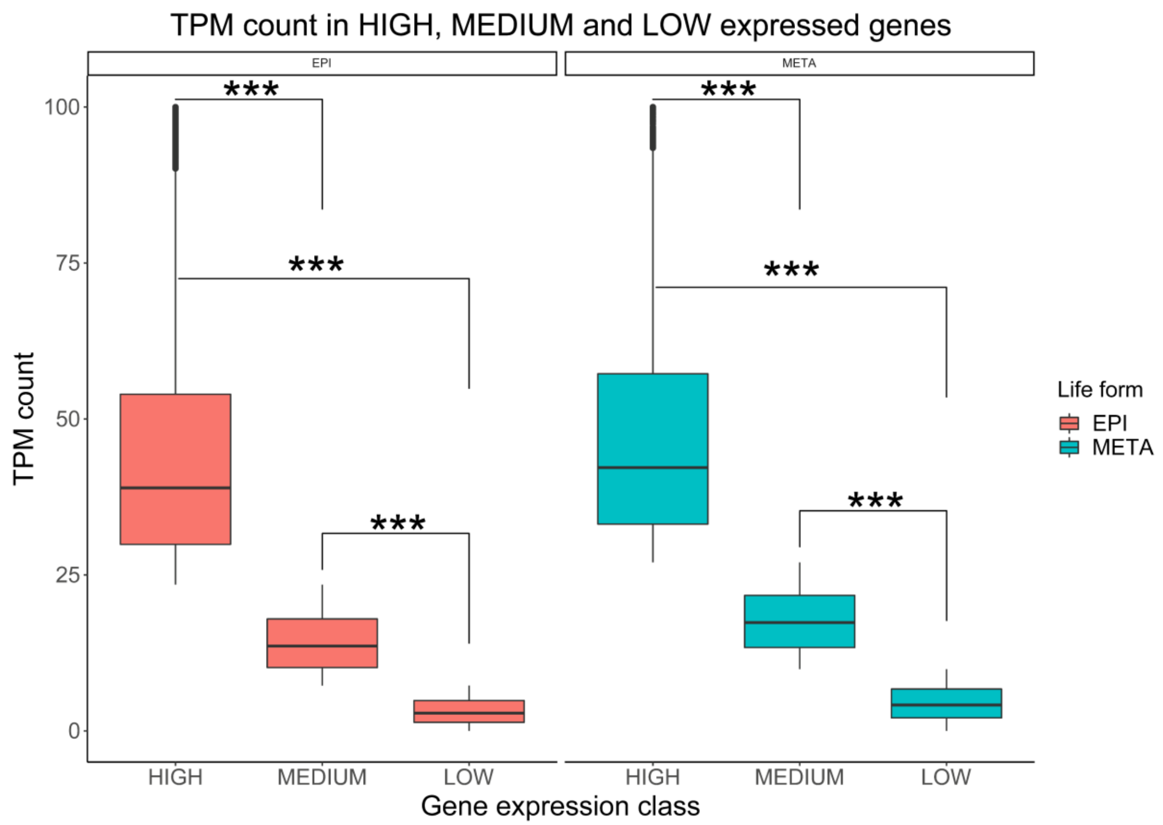


**Figure S13.** Genes classified as high, medium and low expressed based on TPM values for epimastigote (red) and metacyclic trypomastigote (blue) life forms. *** Wilcoxon-Mann-Whitney test with p-value = 0.001.

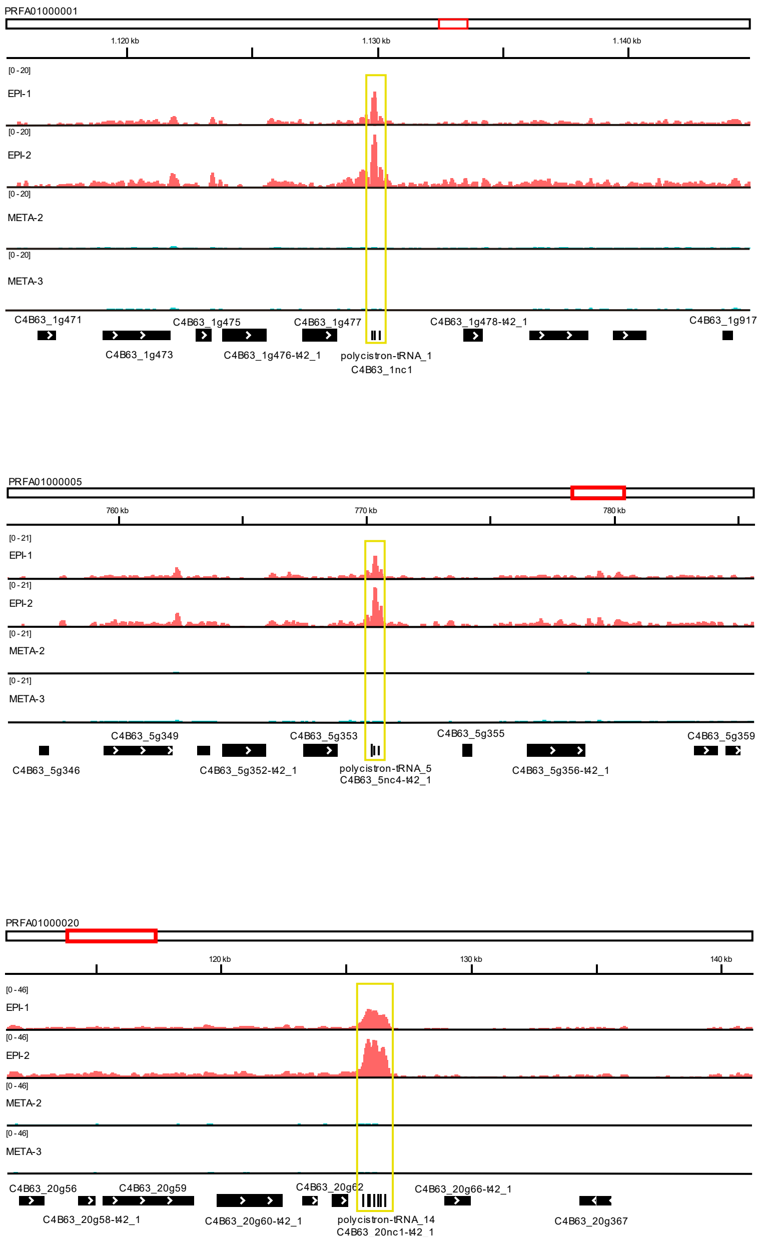


**Figure S14.** IGV snapshots of FAIRE-seq enrichment in epimastigotes (duplicates – red) compared to MTs (duplicates – blue) in three different loci. tDNAs are highlighted in yellow rectangles.

A.

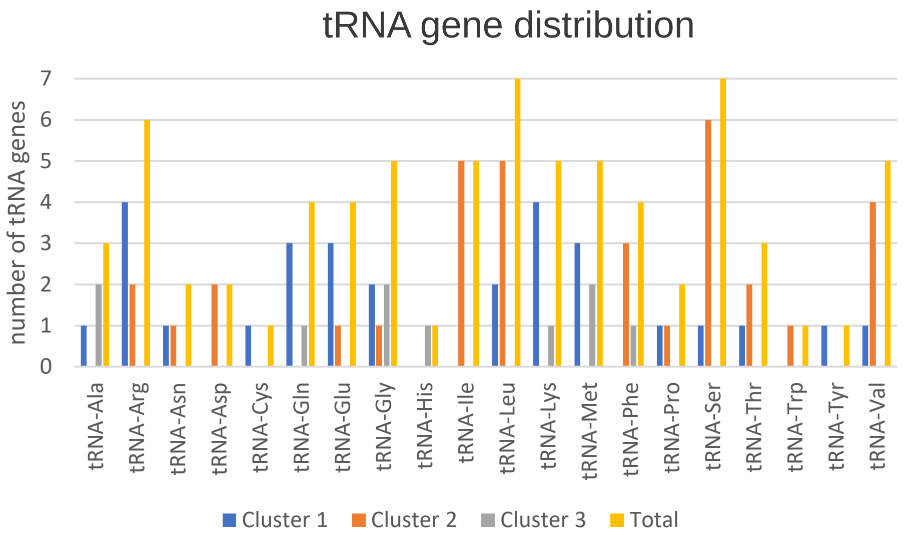


B.

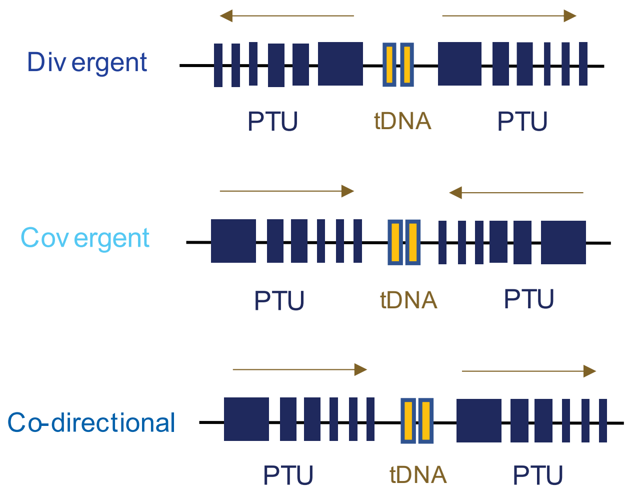


**Figure S15. (A)** Number of tRNA genes in each cluster. The tRNA genes grouped in each cluster after hierarchical clustering, as shown in Figure 6C, were counted and plotted in column graphics according to R group classification of the amino acids. (B) Representative scheme of tRNA gene localization at divergent, convergent and codirectional PTUs. The arrows indicate the sense of transcription in each PTU.

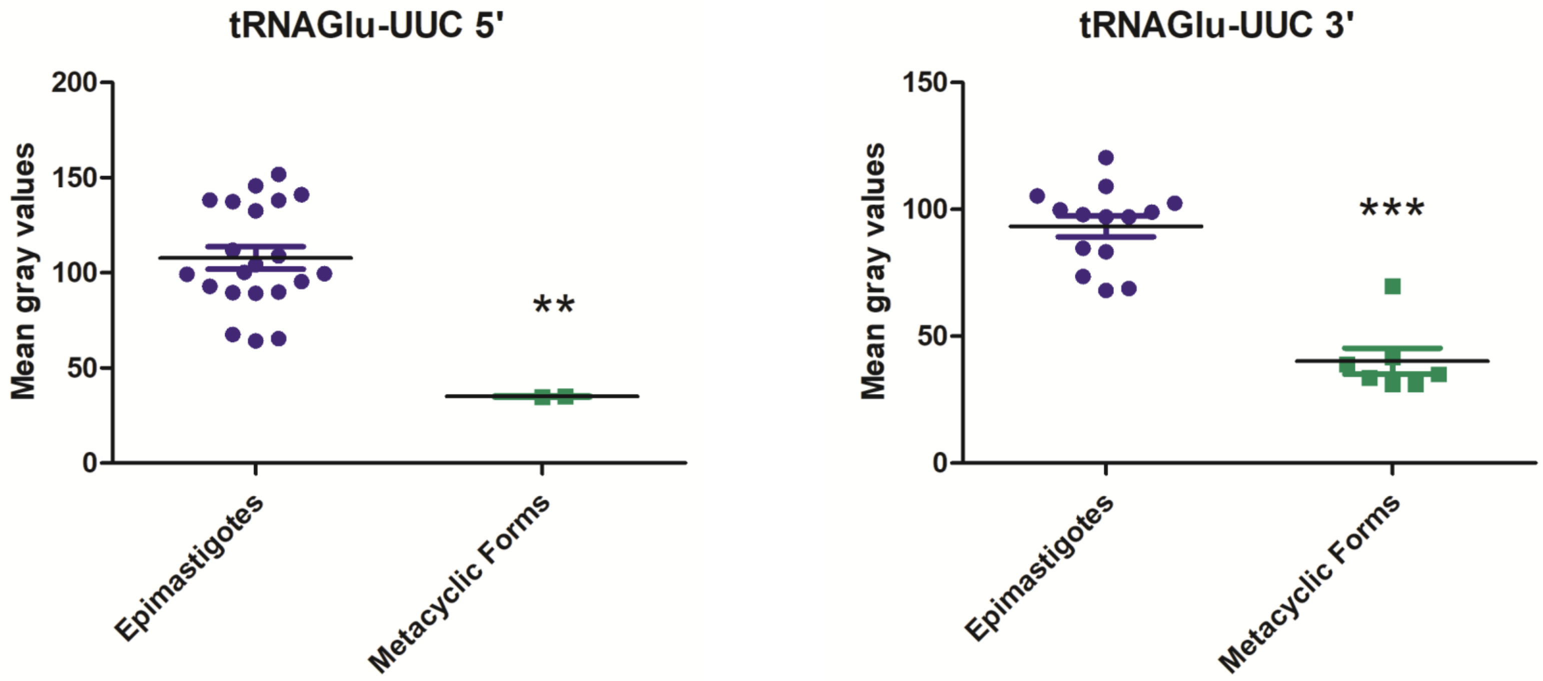


**Figure S16.** Quantification of tRNA-FISH UUC in epimastigote (blue) and MT (green) forms. Mean gray values represent intensity per area. Unpaired T-test: ** (p-value > 0.05), and *** (p-value < 0.0001).
